# Neural evidence for recognition of naturalistic videos in monkey hippocampus

**DOI:** 10.1101/2020.01.03.894394

**Authors:** John J. Sakon, Wendy A. Suzuki

## Abstract

The role of the hippocampus in recognition memory has long been a source of debate. Tasks used to study recognition that typically require an explicit probe, where the participant must make a response to prove they remember, yield mixed results on hippocampal involvement. Here, we tasked monkeys to freely view naturalistic videos, and only tested their memory via looking times for two separate novel v. repeat video conditions on each trial. Notably, a large proportion of hippocampal neurons differentiated these videos via changes in firing rates time-locked to the duration of their presentation on screen, and not during the delay period between them as would be expected for working memory. Single neurons often contributed to both retrieval conditions, and did so across many trials with trial-unique video content, suggesting they detect familiarity. The majority of neurons contributing to the classifier showed an enhancement in firing rate on repeat compared to novel videos, a pattern which has not previously been shown in hippocampus. These results suggest the hippocampus contributes to recognition memory via familiarity during free-viewing.

**Significance Statement:** Recognition memory enables distinction of new from previously encountered stimuli. In the majority of recognition memory work, humans or animals are tasked to explicitly identify whether a stimulus is old or new. In some of these studies—but for unknown reasons not others—there is evidence of hippocampal involvement. Here, when we present trial-unique, naturalistic videos to monkeys, firing rates from many single hippocampal neurons surprisingly differentiate new from repeated videos. The ability of these hippocampal neurons to detect recently-viewed stimuli, despite the videos’ unfamiliar content, suggests they contribute to memory. These results are consistent with hippocampal involvement in the familiarity aspect of recognition memory during free-viewing.

## Introduction

The ability to identify a previously experienced stimulus is termed recognition memory (1). There is debate to what degree the hippocampus—a brain region typically linked to associative and episodic memory (2)—contributes to recognition memory (3). That is, does the hippocampus contribute beyond its role in recollection, or is familiarity, the sense of recognizing a stimulus without recalling specific details, achieved by extrahippocampal regions without hippocampal support (4, 5)? Recent reviews, in light of mixed evidence from lesion work in monkeys and amnesic studies in humans, suggest that the hippocampus may be involved in the familiarity component of recognition—in absence of recollection— depending on task parameters like stimulus type and frequency of stimulus repetition during and across sessions (3, 6). One specific prediction is that the hippocampus only supports recognition memory for stimuli repeatedly used within or across sessions (3).

Neurophysiological work in humans has found hippocampal neurons that differentially respond to repeat v. novel images (7-11), with some studies showing evidence the hippocampus contributes to recognition memory via both recollection and familiarity (9, 10, 12). The neural signature of recognition in this work is typically repetition suppression (13), with decreased responses on repeat compared to novel presentations (10, 11, 14, 15), similar to the neural responses to repeated stimuli in extrahippocampal regions that are more commonly associated with familiarity (13, 16). Notably, these recognition memory studies almost exclusively use designs with explicit memory probes (i.e. old/new judgments) (7-11), even though the familiarity component of recognition memory is more closely tied to implicit memory processes (17). Meanwhile, numerous studies with similar explicit task designs in monkeys have reported little or no responses from single hippocampal units (1, 18-21).

Why is evidence for hippocampal involvement in recognition memory so varied? Recent studies in monkeys (22, 23), which unlike previous monkey work found a large proportion of hippocampal neurons that differentiated repeat from novel images, suggest two possibilities. First, unlike in the aforementioned human and monkey work, animals in these tasks were not required to make an explicit response to probe their memory. This task design aligns with work in humans showing explicit memory probes—as opposed to incidental probes—cloud the neural responses in hippocampus (24, 25), possibly because the requirement of a memory decision recruits additional brain regions that support strategy (e.g. recall-to-reject (25)) or integration of evidence from memory (26). A second difference in these studies is animals had volitional control of image presentation, as images were removed when their eye gaze left the image (22, 23). Hippocampal activity has been linked to both saccades (27, 28) and scanpaths (29, 30), making it unknown if these previous works reflect a brief surge in hippocampal activity after repeat presentation or are more universal familiarity signals maintained throughout the course of longer repeat presentation.

To better understand hippocampal involvement in recognition memory, we recorded single neurons from monkey hippocampus as they engaged in a task with two separate memory conditions on each trial (Fig. 1A), but with no explicit probe or animal control of video presentation. Monkeys freely viewed 6 s episodes (“**clips**”), comprised of three, contiguous 2 s videos (the 2 s videos are designated a, b, and c), sampled from Internet videos of people, animals, animations, and other such engaging content. In each trial, after three presentations of the 6 s clips (the encoding phase), the animals were then shown two, 2 s “**subclips**” with a 1 s delay period in between (the retrieval phase). In subclip 1, either the first 2 s of the original clip was shown (repeat 1), or a new 2 s subclip was shown (novel 1). Following the 1 sec delay period, subclip 2 could be either the last 2 s of the original clip (repeat 2), or a new 2 s subclip (novel 2). Reward was separated from the memory component as juice reward was given for achieving fixation before each clip and retrieval period, and a second fixation was required in each case to proceed with video presentation (Fig. 1A). Crucially, once a clip or subclip was used in a given trial, it was never again used for that monkey, thereby making the novel videos a true first presentation, and the repeat videos targeted probes of memory for recently-shown videos.

**Fig. 1.**
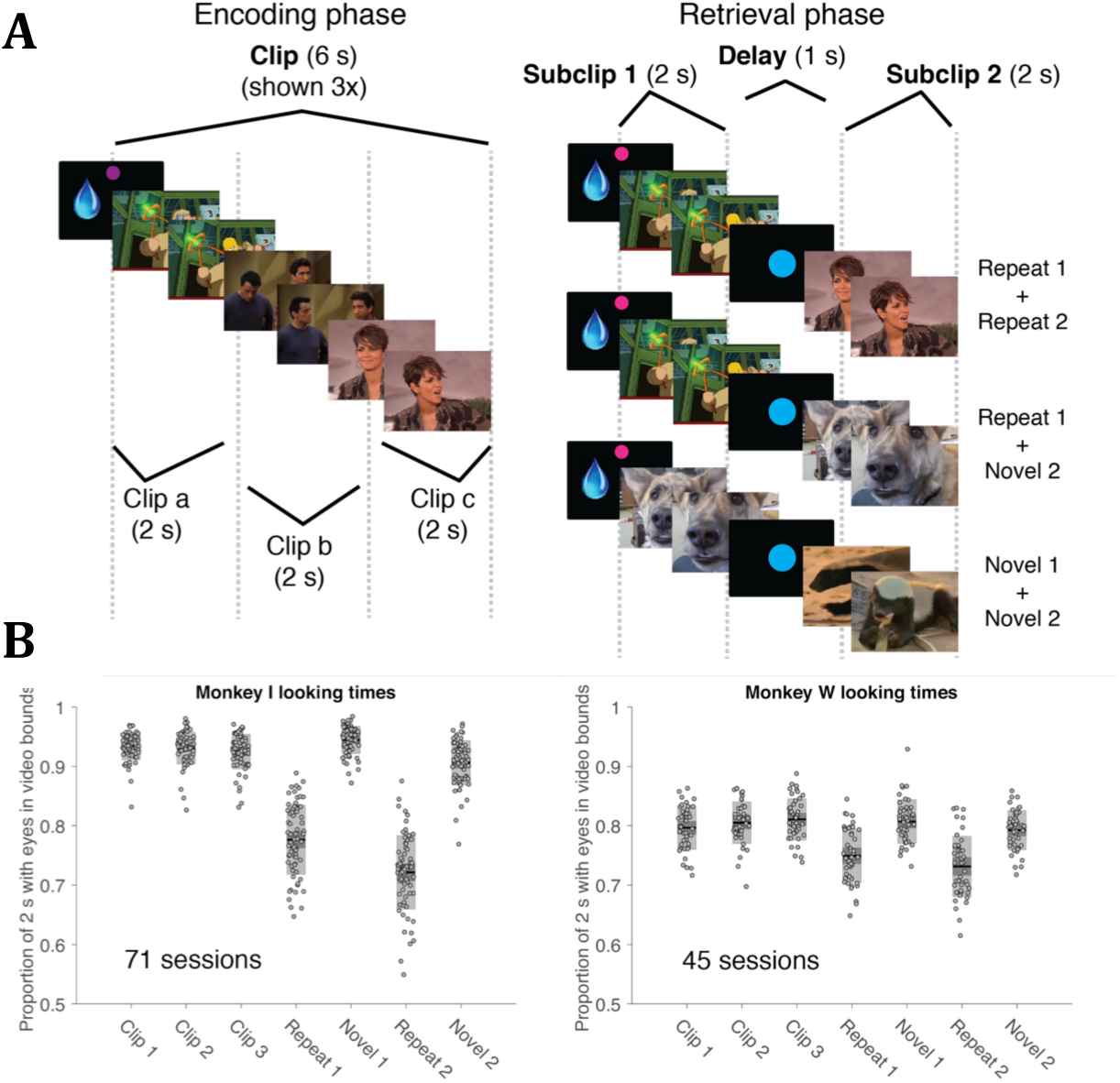
Task design and behavior. **A)** In each trial, monkeys were shown 6 s clips, which were comprised of three, 2 s movies (sec a, b, and c) spliced together from random video sources (Metho Each clip was repeated 3x during the encoding phase. In the retrieval phase, monkeys were shown two, 2 s subclips surrounding a 1 s delay (a delay of only 1 s was used to maintain monkey attention). Three trial types were equally likely during this pha subclip 1 is a repeat of the first 2 s of the clips and subclip 2 is a repeat of the last 2 s of the clips, ii) subclip 1 is a repeat of the first 2 s of the clips and subclip 2 is a novel video, and iii) subclip 1 and subclip 2 are both novel videos. Each liquid drop icon indicates fixation “game” where animals were given juice reward for achieving 0.5 s fixation to a dot. After reward was given, monkeys had to r acquire fixation for 0.5 s to begin the upcoming video. Colors of fixation dots are as shown, with a purple dot indicating an upco clip and a pink dot denoting entry into the retrieval phase. **B)** T proportion of time each monkey’s eye gazes were within the bounds of the movie for section a of clips 1-3, and for each of the 2 s subclip periods, averaged over sessions. Note that Repeat 1 (shown during subclip 1) is a repeat of the first 2 s of the clips (section a), while Repeat 2 (shown during subclip 2) is a repeat of the last 2 s of the clips (section c). Therefore, the middle 2 s of the clip (section b) never shown during retrieval. Shaded boxes are 95% confidence intervals (dark gray) and the SDs (light gray).

Hippocampal responses to repeated videos in this task address a number of the questions outlined above concerning the hippocampus’s involvement in recognition memory. A large proportion of single neurons significantly changed their firing rates during both repeat v. novel conditions (subclip 1 and subclip 2), and did so consistently across trials despite the use of trial-unique content within and between sessions. Surprisingly, these neurons showed a predominantly repetition *enhancement* in firing rates, which was also time-locked to the 2 s duration of each subclip period. Many of the same neurons showed this pattern of firing for both the subclip 1 and subclip 2 periods, despite not continuing for the delay period between them, suggesting hippocampal neurons support familiarity detection limited to the duration of stimulus presentation.

## Results

### Behavior

To determine if monkeys engaged with the task, we looked for evidence of each animal remembering videos during each trial via a preferential looking paradigm, in which memory is associated with viewing repeated stimuli for shorter times than novel stimuli (22, 23, 31, 32). We measured the time each monkey looked within the bounds of the video stimuli during the first 2 s of all three clips, as well as during the two, 2 s subclip periods for 116 sessions (Fig. 1A; 71 monkey I and 45 monkey W). The looking times for each of these periods are averaged for each session and then the proportion of the 2 s the animal looked at the video is plotted separately for each monkey (Fig. 1B). We anticipate that for both subclip 1 and subclip 2, when monkeys are shown a repeat instead of a novel video, on average they should look for less time than novels (note that, when either subclip is a repeat, it is the 4^th^ presentation of that video, since all repeat videos were also shown during clips 1-3). Indeed, for each monkey, both during the subclip 1 (Monkey I, p=5.8e-24; Monkey W, p=2.4e-7; Wilcoxon rank-sum test, FWE-corrected) and the subclip 2 (Monkey I, p=2.1e-23; Monkey W, p=2.4e-7; Wilcoxon rank-sum test, FWR-corrected) periods, monkeys looked at repeat videos for significantly less time than novel videos. Each monkey also looked at the novel videos during the subclip 1 period slightly longer than clip 1 (Monkey I, 0.933 for clip 1a v. 0.945 for novel subclip 1, p=0.0049; Monkey W, .797 for clip 1a v. .807 for novel subclip 1, p=1.0; Wilcoxon rank-sum test, FWE-corrected), although monkey I’s preference for novel subclip 1 was not significantly greater than monkey W’s (p=0.94, interaction in mixed effects linear regression). The significant differences in looking times between novel and repeat videos, in addition to the high proportion of time monkeys looked at novel videos (>0.90 for monkey I and >0.79 for monkey W for all novel clips and subclips), indicate monkeys were engaged in the task.

To quantify how well monkeys classify repeat v. novel videos in our task, we used looking times in a signal detection theory framework similar to a recent publication (23). Our expectation is that since monkeys on average look longer at novel videos than repeat videos, by comparing their clip 1 looking times (a novel video is always shown during clip 1) to the subclip 1 looking times, we can assess the rates at which the monkeys identify novel or repeat videos shown during subclip 1. Since on average when including both novel and repeat videos monkeys looked at the subclip periods for shorter times than the clip periods (Fig. 1B), we used a constant threshold of 0.8 to segregate our behavior (Methods). That is, if the monkeys look at a novel video in subclip 1 greater than 80% of the time they looked at clip 1a (the first 2 s of clip 1), we consider that a correctly identified novel (true positive). And if the monkeys look at a repeat video in subclip 1 greater than 80% of the time they looked at clip 1a, we consider it incorrectly identified (i.e., false positive). When we apply this process to the 116 sessions across both monkeys (average of 125.9±2.5 trials per session), on average for the 1^st^ subclip we can classify 85.0±0.9% of novel videos as novel compared to 66.4±0.8% of repeat videos as novel, which yields an average across sessions of d’=0.70±0.05 (SE for all errors). In addition, when we use the same process to compare the looking times from the novel/repeat videos from the 2^nd^ subclip v. the last 2 s of the clip 1 period (clip 1c), we can classify 80.2±0.9% of novel videos as novel compared to 54.1±1.0% of repeat videos as novel, which yields d’=0.80±0.05. Classification was at similar levels when using clip 3 to compare to the subclips instead of clip 1 (clip 3a v. subclip 1: d’=0.72±0.05; clip 3c v. subclip 2: d’=0.63±0.04). We find weaker, but still well above chance, classification using a threshold of 1.0, as is typically used in preferential looking (Supplementary text). This strong classification of the subclip periods gives us additional behavioral evidence that the monkeys recognize videos in both subclip periods that were recently shown during the clip periods.

### Single neuron evidence of recognition memory

We recorded from 209 single units in the hippocampus (SI Appendix, Fig. S4) of the same two monkeys from our behavioral analysis during 116 sessions (unit breakdown: Rhesus I = 120 units, Rhesus W = 89 units). In the limited previous work that has used naturalistic video stimuli during electrophysiological or neurophysiological recordings—all in humans—neurons are responsive throughout the medial temporal lobe (33) and hippocampus (34, 35). Therefore, while we limited our recordings to hippocampus to investigate our main questions about its involvement in recognition memory, we did not attempt to limit our recordings to a particular subfield (note that during such deep brain acute recordings localization is limited to about ±1 mm in all three dimensions, which would make the subfield within hippocampus uncertain for many units). The firing rates of the neural population during video presentation were variable [mean firing rate ± SD for 155 neurons <10 Hz (36) = 2.8 ± 2.6 Hz; mean firing rate ± SD for 55 putative interneurons ≥10 Hz = 26.6 ± 16.8 Hz] but with a similar breakdown as our previous work recording hippocampal neurons during serial image presentation (23). The population sparseness (37) of neural firing to novel and repeat videos was *a*^P^ = 0.30, similar but slightly more sparse than the *a*^P^ = 0.33 (37) and *a*^P^ = 0.37 (23) found in previous studies of monkey hippocampal firing rates to images.

Importantly, we made no effort to select for stimulus-responsive neurons, but instead during daily recordings lowered the electrode into hippocampus and recorded from the first stable single unit(s) we could isolate. Monkeys were head-fixed during the initial isolation process and rested in front of a black computer screen. Therefore, our hippocampal population was not pre-selected in any way for visual or video stimulus responsiveness.

We predicted our task might elicit recall in monkeys, which we hoped to detect during the delay period by cueing with the repeated videos shown in subclip 1. This design was inspired by work showing single hippocampal neurons respond to free recall of recently-viewed video stimuli in humans (33). We found no evidence for such cued recall signals (SI Appendix, Supplementary text & Figs. S1-S2), with hippocampal neurons typically returning to basal firing levels during the delay or ITI periods after repeat presentation as shown in both our population classifier (Figs. 2-3) and single unit examples (Fig. 4; SI Appendix, S5). However, we found strong hippocampal changes in firing rate to repeat v. novel stimuli throughout their presentation during the subclip 1 and subclip 2 periods. As we address in the discussion, we believe these results are best explained by the hippocampus supporting the familiarity component of recognition memory.

**Fig. 2.**
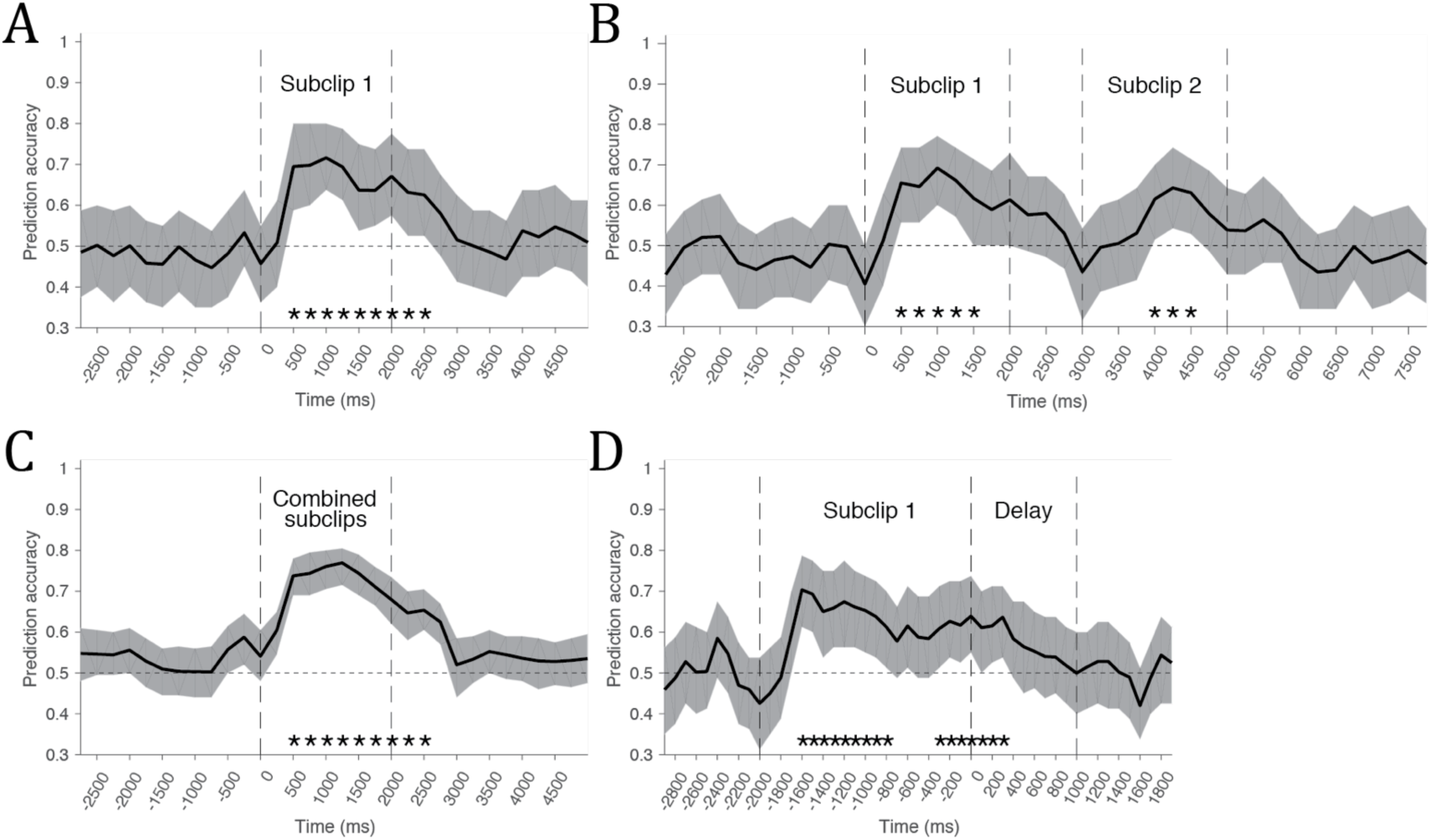
Classification of novel and repeat videos via firing rates of neural population. **A)** Prediction accuracy of novel v. repeat video identification aligned to subclip 1 onset for all trials combined. Dashed vertical lines indicate subclip 1. **B)** Prediction accuracy when classifying trials where the monkey was shown either novel+novel (1/3^rd^ of trials) or repeat+repeat (1/3^rd^ of trials) videos for subclip 1+subclip 2. Dashed lines outline the contiguous subclip 1 (2 s), delay (1 s), and subclip 2 (2 s) periods. **C)** Prediction accuracy when classifying pooled subclip 1 and subclip 2 trials based on whether novel or repeat videos were shown during that period. **D)** Prediction accuracy aligned to delay onset with trials classified on whether novel or repeat videos were shown during the preceding subclip 1 period. Spikes are integrated in windows of 500 ms with a moving window of 250 ms for each point in the first 3 graphs, and integrated for 200 ms with 100 ms steps for the last graph to show higher resolution of the drop-off after delay onset. Error bars for all graphs are the 5^th^ and 95^th^ percentile of prediction accuracy across 200 data subsamples, while adjacent asterisks represent significant clusters of classification above chance from 1000 permutations (Methods).

**Fig. 4.**
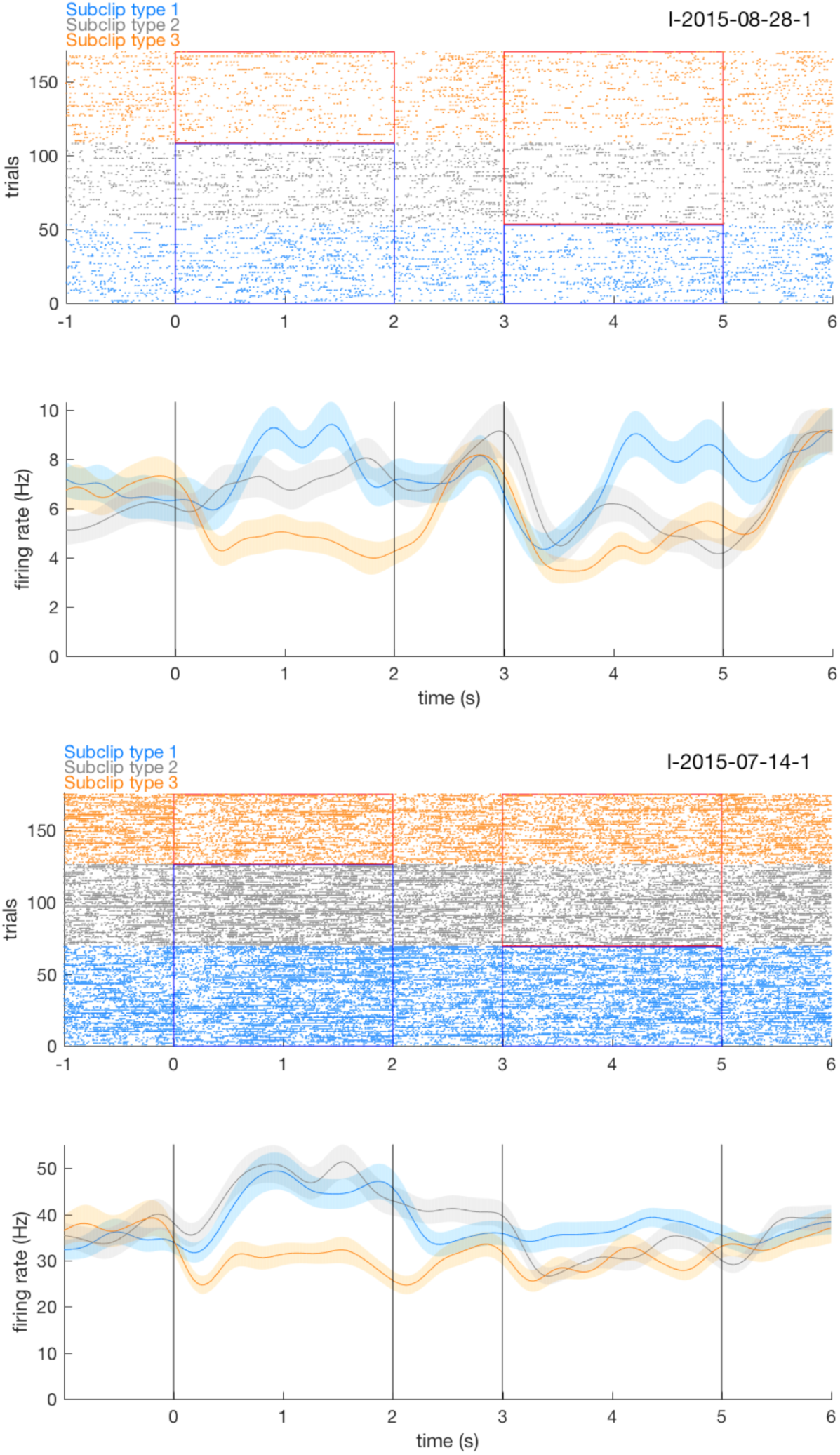
Example neurons. Rasters (top) and probability density functions (PDFs, bottom) for two different neurons. Trials are bro up by subclip type (1, blue: repeat/repeat; 2, gray: repeat/novel; a 3, orange: novel/novel from bottom for subclip 1/2) and aligned to subclip 1 on. Vertical black lines are boundaries of subclip 1 (2 s), delay (1 s), and subclip 2 (2 s). In the rasters, areas outlined in red indicate presentation of novel videos, while areas outlined in blue indicate repeat videos. For example, subclip type 2 trials in gray a when monkeys are shown a repeat video for subclip 1 and after the. delay a novel video for subclip 2. Dark lines in the PDF are average firing rate and outlines are SE

The subclip periods act as the key recognition memory conditions in our task, as every subclip 1 had a 2/3^rd^ chance of a repeat video and every subclip 2 had a 1/3^rd^ chance of a repeat video. First, we asked if hippocampal neurons changed their firing rates during novel or repeat video presentation as compared to baseline firing rates during fixation periods. Indeed, we found approximately half of neurons were significantly modulated by either novel or repeat video presentation during either subclip period, which is comparable to previous breakdowns of neurons in monkey hippocampus during presentation of novel or repeat images (Table 1, “Total video responsive”). However, the proportion of neurons that increased in firing rates to repeat videos, with 44% increasing compared to only 20% decreasing during subclip 1 (p=6.8e-5, Χ^2^-test, FWE-corrected), and 31% increasing compared to only 13% decreasing during subclip 2 (p=3.0e-4, Χ^2^-test, FWE-corrected), were both larger than expected by chance. This greater number of neurons increasing rather than decreasing in firing rate to repeat videos is surprising, considering 1) that repeated stimuli tend to universally show repetition suppression of neural responses in medial temporal lobe (13, 14) and 2) previous single unit responses in monkey (22, 23) and human (11) hippocampus in particular have shown predominantly suppressed firing rates when participants are shown repeat images (Table 1, last 2 columns). We considered it possible that while novel and repeat firing rates were enhanced compared to baseline (fixation period firing rates), they might actually be suppressed compared to responses during clip 1 (essentially the first novel video on each trial). If this were true, we’d expect clip 1 firing rates to be higher than baseline. Instead, for most neurons clip 1 firing rates were lower than baseline, and particularly so for those neurons showing the strongest repetition enhancement signal (SI Appendix, Supplementary text & Figs. S3).

**Table 1.**
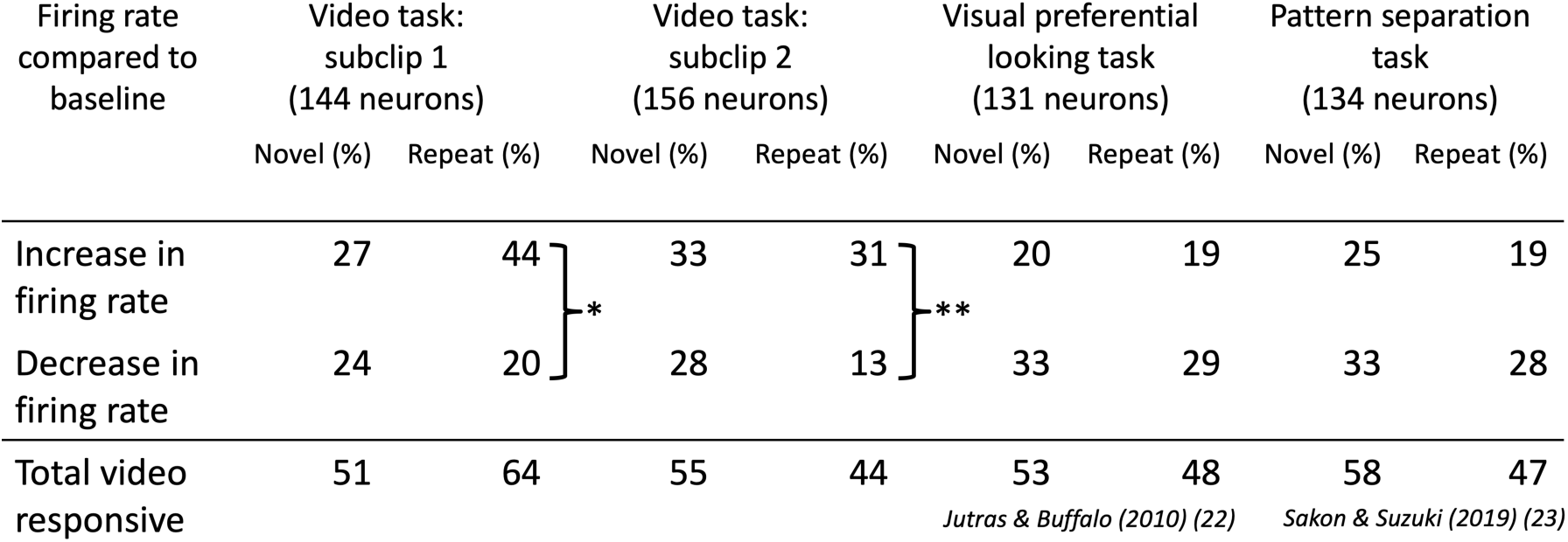
Comparison of stimulus responsive neurons between video task and image recognition tasks. The percentage of monkey hippocampal neurons that significantly increase or decrease their firing rates from baseline (fixation period firing rate) for the subclip periods of the video task (p<0.05, Wilcoxon rank-sum) as well as image presentation in two previously published recognition memory tasks (22, 23). The bottom row is the total percentage of stimulus responsive neurons (the sum of the first two rows). *p=6.8e-5, **p=3.0e-4, FWE-corrected.

Next, we directly compared neural firing rates between repeat v. novel videos (as opposed to comparison to baseline firing rates). During the subclip 1 period, 27/144 (19%) neurons fired significantly more to repeat than novel videos (p<0.05, rank-sum test), as compared to only 7/144 (5%) where neurons fired more to novel than repeat videos. These differences in proportions are not expected by chance (p=1.0e-3, Χ^2^-test, FWE-corrected). Similarly, during the subclip 2 period, 21/156 (13%) neurons fired significantly stronger to repeat than novel videos, as compared to 8/156 (5%) where neurons fired more to novel than repeat videos, which again is not expected by chance (p=0.04, Χ^2^-test, FWE-corrected). Once again, these proportions of repetition enhancement neurons were greater than what has been shown in hippocampal neurons recorded while monkeys performed a serial image task in our previous work (23), where 12/134 (9%) neurons showed significantly greater firing rates to repeat than novel images, while 15/134 (11%) showed significantly greater firing rates to novel than repeat images. Therefore, unlike in the serial image task, hippocampal neurons recorded during the video task respond in a manner more consistent with repetition enhancement.

We made similar comparisons after using our behavioral measure to classify trials, wherein we only compare firing rates for repeat videos when looking times indicated the monkey recognized the repeat v. novel videos when the looking times indicated the monkey recognized the video as novel. For these behaviorally-identified trials, during the subclip 1 period, 38/152 (25%) neurons fired significantly stronger to repeat than novel videos, as compared to only 9/152 (6%) where neurons fired more to novel than repeat videos (p=8.4e-6, Χ^2^-test, FWE-corrected). During the subclip 2 period, 33/147 (22%) neurons fired significantly stronger to repeat than novel videos, as compared to only 3/147 (2%) where neurons fired more to novel than repeat videos (p=1.9e-7, Χ^2^-test, FWE-corrected). Notably, of the 38 neurons with higher firing rates to repeat videos in subclip 1 and the 33 neurons with higher firing rates to repeat videos in subclip 2, 18 of these neurons are the same, indicating these neurons are tuned to repeat video detection regardless of the period. Note that different numbers of neurons appear in the denominator for each of the periods tested above as a minimum firing rate threshold and number of trials recorded for each period had to be met for a neuron to be included in that test (Methods).

### Population evidence of recognition memory

Considering the strong, differentiable responses at the individual neuron level to novel and repeat videos, we asked if we could classify novel and repeat videos in the subclip periods via the firing rates of our neural population. First, focusing on subclip 1, we took the 110 neurons with a minimum firing rate of 0.5 Hz (69 neurons <10 Hz = 2.6 ± 2.5 Hz; 41 putative interneurons ≥10 Hz = 25.2 ± 20.1 Hz) that were recorded for at least 35 novel and 35 repeat trials and created a 10-fold, cross-validated logistic regression classifier (23, 38) on this pseudopopulation. This classifier trains each fold on 90% of trials and tests on the held out 10% in a balanced design, thereby using every trial once in one of the ten test sets. Since we use the neural responses to predict a binary classification, we are only testing if hippocampal neurons generally categorize novel v. repeat videos agnostic to the content of the videos themselves. We trained separate classifiers for spikes integrated over each 500 ms time window that was stepped through in 250 ms increments locked onto stimulus presentation. As most neurons in the pseudopopulation were recorded for more than 35 novel or repeat trials (mean trials ± SE: 79.8 ± 1.5 novel and 77.5 ± 1.3 repeats), we ran each classifier for 200 permutations and randomly subsampled from the total trials recorded for each neuron. The average prediction accuracy from these permutations is shown in Fig. 2, with significant clusters of classification assessed using a maximum statistic method that accounts for multiple comparisons over time (39) (Methods).

The prediction accuracy of novel v. repeat for subclip 1 was significantly above chance from 500-2500 ms after video onset (Fig. 2A), indicating the population of hippocampal neurons can differentiate new from old videos via a rate code. This period of classification matched the 2 s length of subclip 1, albeit with an offset from image presentation of ∼500 ms, close to the 300-400 ms latencies typically shown in human hippocampal recordings to naturalistic stimuli (40).

For the analysis just described, we used all three retrieval phase trial types (Fig. 2A) to train and test the classifier. And since we trained the classifier for subclip 1— where we used the predictor “repeat” for the first two trial types and “novel” only for the third trial type—we cannot simultaneously classify subclip 2 as the predictor would be “repeat” only for the first trial type. Therefore, to use the same classifier across both subclip 1 and subclip 2 simultaneously, we removed the middle trial type, thereby only classifying trials where the monkey was shown either repeat+repeat or novel+novel for subclips 1+2. Despite using only these 2/3^rd^ of total trials, the prediction accuracy of novel v. repeat was significantly above chance for clusters in both the subclip 1 and subclip 2 periods (Fig. 2B). Classification across both periods using the same classifier suggests a similar rate code underlies repeat video recognition in each period.

Next, since the classification for the subclip 1 and subclip 2 periods were both novel v. repeat, we can essentially double our trial count if we pool both periods together and run the same novel v. repeat classifier. In this case, prediction accuracy approaches 80% at its peak, ∼10% higher than the accuracy when looking at only subclip 1, and once again is significant from 500-2500 ms after video presentation (Fig. 2C). Stronger classification when pooling trials across the two subclips further suggests that neurons utilize a similar code for repeat video recognition across the two repeat v. novel conditions.

The preceding classifiers show two distinct periods of novel v. repeat classification during the subclip 1 and subclip 2 periods, while classification does not appear to carry through into the delay period.

However, to more precisely confirm that the identity of novel v. repeat videos is not held by neurons into the delay, we created a classifier that is time-locked to the beginning of the delay period with higher resolution (200 ms integration window, 100 ms steps) to better identify when classification ceases (Fig. 2D). Indeed, the presentation of novel v. repeats in the subclip 1 period were no longer significantly classified within 300 ms of the video finishing, consistent with the 300-400 ms latencies typically shown with such stimuli in hippocampus (40), suggesting that hippocampal neurons do not hold the memory of the previous video into the delay between subclips. The same drop in firing rate upon onset of the delay period can also be seen at the level of individual neurons, with the differences between firing rates to novel and repeat videos quickly disappearing in all six of the example neurons that contribute strongly to the subclip 1 classifier (Figs. 4 and SI Appendix, Fig. S5).

While these analyses (in particular, Fig. 2B-C) imply that the population of hippocampal neurons classify novel and repeat videos via the same rate code both 1) within each 2 s subclip period and 2) between the separate subclip 1 and subclip 2 periods, since we train and test our classifier at each time bin, it’s possible different sets of neurons or the same neurons with different signals over time are responsible for the clusters of classification we see in Fig. 2. To show that the same set of neurons use a stable rate code over time, we can train the classifier on one time bin and then test (cross-validate) the classification on a different time bin (41). This analysis can address 1) similar coding within each subclip, since we can train and test time bins within the same subclip period, and also 2) between the two subclips, since we can train in a subclip 1 time bin and test in a subclip 2 time bin (and vice versa). Once again, as in the analysis for Fig. 2B, we only use the 2/3^rd^ of trials where the monkey saw repeat+repeat or novel+novel for subclips 1+2.

A heatmap of the classification accuracy when we train and test at each time bin shows evidence of a stable neural code both within and between subclips (Fig. 3). Prediction accuracy approaching 70% can be achieved 500 ms into the subclip 1 period (first white box) and shows approximately equivalent accuracy for all combinations of training and testing within the next 1500 ms. A similar phenomenon is present within the subclip 2 period, albeit with prediction accuracies only approaching 65% as in the subclip 2 period of Fig. 2B. Meanwhile, stable regions of prediction accuracy occur if we train on the subclip 1 period and test in subclip 2 or train on the subclip 2 period and test in subclip 1, indicating that the same rate code classifies novel v. repeat videos in both periods.

**Fig. 3.**
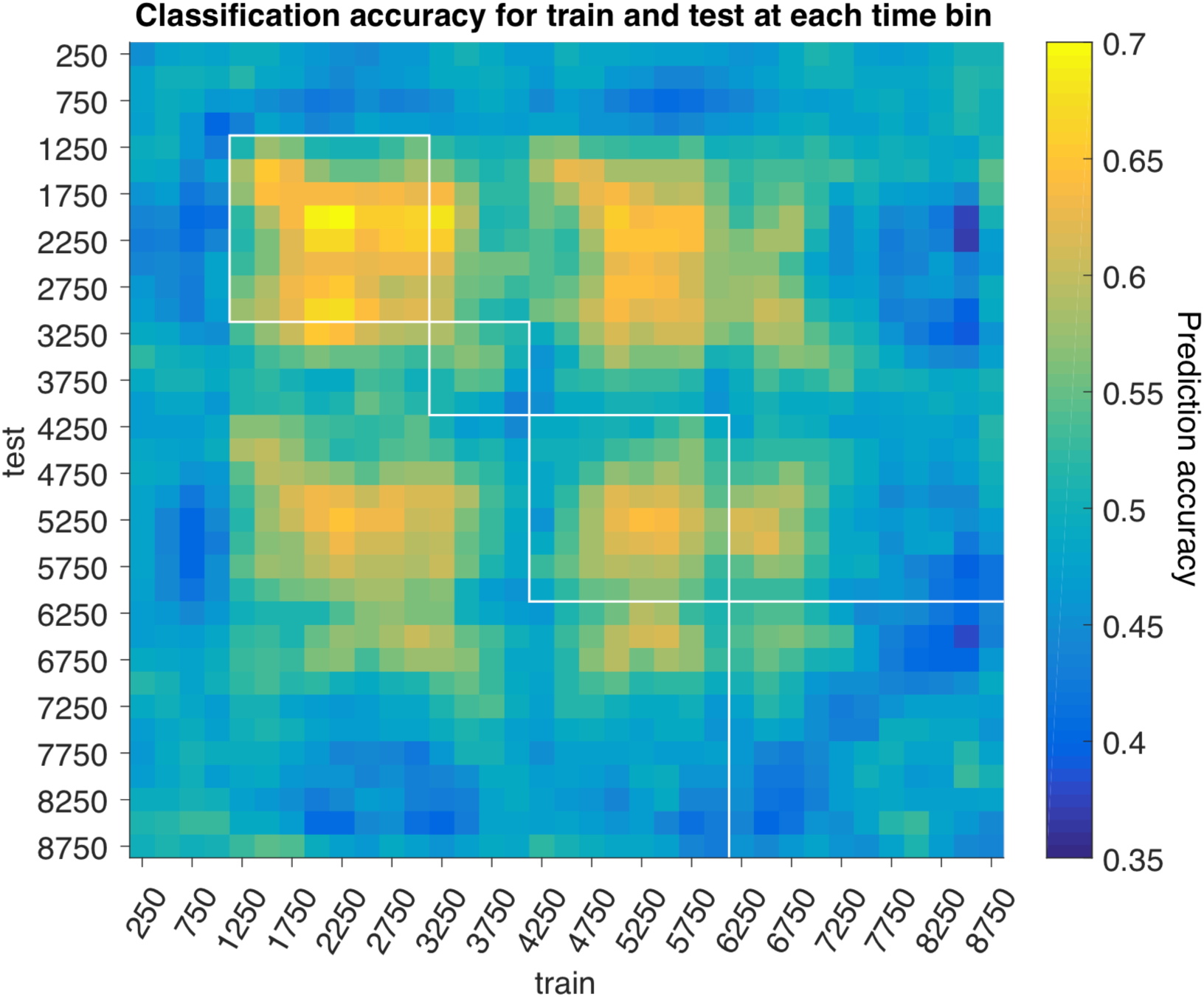
Training and testing classifier at different times reveals a stable neural code. Using the same classifier detailed in Fig. 2, prediction accuracy of novel v. repeat videos was assessed via 10-fold cross-validation at each point in the heatmap by training on one time bin and testing on the other. White boxes—in order left to right—indicate subclip 1, delay, subclip 2, and ITI periods. The diagonal of the heatmap is equivalent to the prediction accuracy in Fig. 2B.

Next, we investigate the individual neurons that contribute most to these classifiers to understand the neural code underlying recognition of novel v. repeat videos. Two example neurons are shown in Fig. 4 (4 additional examples are shown in SI Appendix, Fig. S5, along with encoding period rasters for all six example neurons in SI Appendix, Fig. S6), which represent the neurons with the 2^nd^ and 12^th^ most positive weights in the classifier for the subclip 1 period and the 5^th^ and 12^th^ most positive weights in the classifier for the subclip 2 period (Fig. 2). In our classifier, positive weights are indicative of classification via higher firing rates to repeat than novel videos. Therefore, negative weights would indicate higher firing rates to novel than repeat videos, while weights of 0 mean no difference in firing rates between novel and repeat videos. The distribution of weights across the population for the classifiers for the two subclip periods—after setting the first neuron’s weight to 1 and normalizing the remaining neurons based on this value—are shown in Fig. 5A-B (red).

**Fig. 5.**
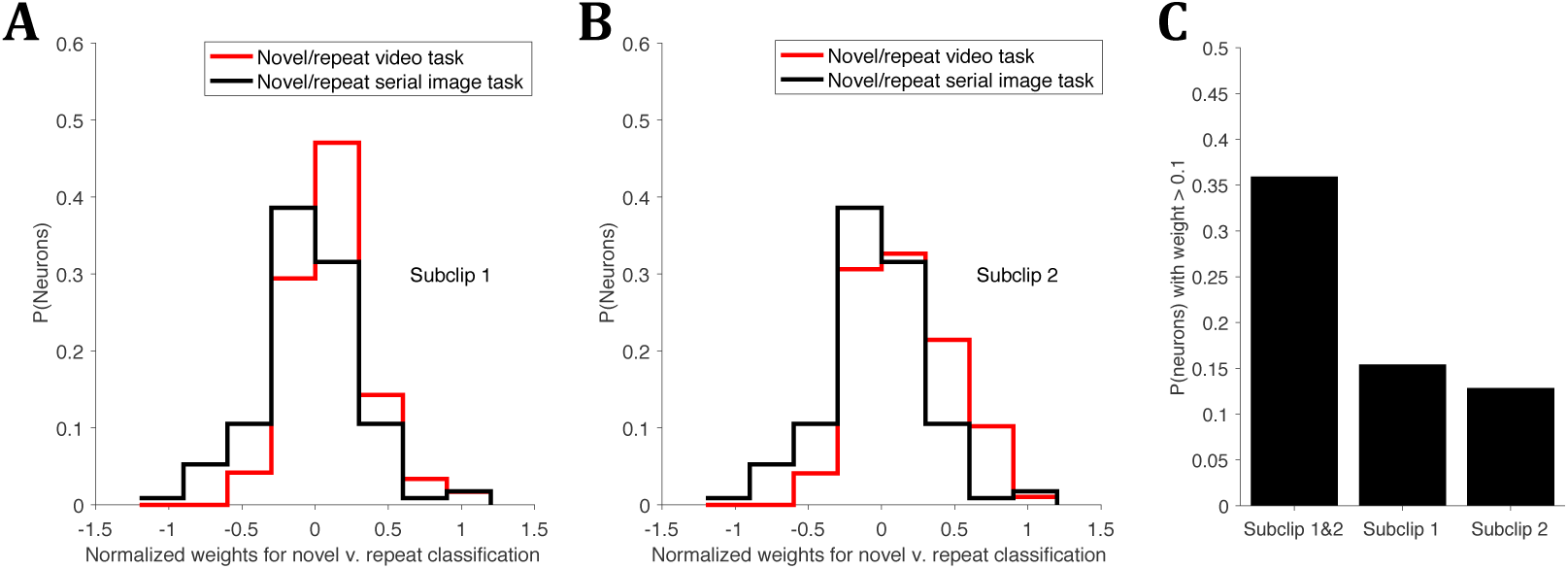
Distribution of weights contributing to population classifier. **A)** The red line shows the neurons for the classifier trained and tested on subclip 1 as in Fig. 2A, but for a single time bin across the subclip 1 period. The black line is for the same classifier trained and tested on hippocampal neurons recorded during a task where monkeys viewed serially presented novel and repeat images (23). The weights for each neuron were ranked and normalized such that the most positive weight in each distribution was set to 1. The weights shown in red here are plotted by anatomical location in SI Appendix, Fig. S4, which shows the neurons that contribute strongly to the classifier are found throughout hippocampus. **B)** Similar to A, but comparing the same classifier trained and tested on the subclip 2 period with the serial image task. **C)** The proportion of neurons with weights for subclip 1 and subclip 2 that were >0.1 (i.e. classifies at least as well as 10% of the best classifying neuron), using the classifiers for subclip 1 and subclip 2 shown in A and B. Subclip 1&2 indicates the weight for those neurons was >0.1 for both periods, while neurons with weights >0.1 for only one of the periods is shown in the next two bars.

The majority of neurons that contribute to the classifier have positive weights, with firing rates to repeat videos greater than novel videos as in the examples of Fig. 4. In fact, when we train a classifier across the entire subclip 1 period (from 500-2500 ms after video presentation), of the 119 neurons in this population, 23 had weights greater than 0.3, while only 5 had weights less than -0.3. This distribution is in stark contrast to our previous hippocampal recordings in monkeys during a serial image task, where animals viewed sequences of images that contained novel, repeat, and lure (similar) images. Plotting normalized weights from the classifier in that task, in which positive also indicates repeat > novel firing rates and negative indicates novel > repeat firing rates, we see a more even distribution of positive and negative weights (Fig. 5A, black). In that case, of the 114 neurons in the population, 15 had weights greater than 0.3, while 19 had weights less than 0.3. The difference between these two distributions is significant (p=0.0013, Wilcoxon rank-sum test). We repeated this analysis for the subclip 2 period from 500-2500 ms after video presentation, and show a similar result (Fig. 5B, p=1.1E-4, Wilcoxon rank-sum test).

In sum, while our previous work (23) (Fig. 5A, black) and others (10, 22, 42) have shown a similar mixture of neurons that code for novel v. repeat recognition via either novel>repeat or repeat>novel firing rates, in our video task hippocampal neurons predominantly show this latter code (repeat>novel firing rates) as in the examples in Fig. 4.

Finally, we return to the question of whether these hippocampal neurons show a general recognition memory response across both subclip periods of the task. In other words, are the same neurons responsible for repeat v. novel video classification in both subclips, or do neurons differentiate these two “contexts” by classifying within only one subclip? Earlier, we found that of the 38 neurons with significantly higher firing rates to repeat than novel videos in subclip 1 and the 33 neurons with significantly higher firing rates to repeat than novel videos in subclip 2, 18 of these neurons are the same. Now, with the use of the classifier, instead of relying on multiple significance tests to group neurons, we can assess the ubiquity of each neuron’s recognition memory response by comparing its weights to the separate classifiers trained on subclip 1 and subclip 2 shown in Fig. 5A-B. Indeed, of the neurons with weights >0.1 in at least one of the periods, 56% had weights >0.1 for both periods (Fig 5C), indicating higher firing rates to repeat than novel videos. Meanwhile, only 24 and 20% of neurons showed weights >0.1 for the subclip 1 and subclip 2 periods, respectively. Therefore, the majority of recognition memory responses—shown via higher firing rates to repeat than novel videos—are general across contexts.

## Discussion

We designed a memory task that, to the best of our knowledge, is the first to utilize naturalistic video stimuli in monkeys while recording from single hippocampal neurons. The animals were engaged in the task, as indicated by significantly greater looking times for novel than repeat videos, a proxy for memory of video content.

Crucially, a large proportion of hippocampal neurons discriminated the presentation of novel v. repeat videos during the two retrieval (subclip) conditions despite there being no explicit probe to test their memory (and therefore no reward linked to successful retrieval). Further, these neurons detected repeated videos during these periods throughout each session even though videos were unique to each trial. The dominant signal for repetition was surprisingly an enhancement in firing rates. Most neurons that showed this repetition enhancement signal in the first retrieval condition also showed the same signal in the second retrieval condition—but it was not held during the delay—suggesting they support familiarity. These results suggest the hippocampus is involved in recognition memory for trial-unique, freely-viewed naturalistic videos.

The involvement of the hippocampus in recognition memory has been hotly debated, with a recent proposal trying to reconcile the mixed literature by arguing the hippocampus is recruited in recognition memory when stimuli are repeatedly used within or across experiments (3). This theory was largely based on numerous studies showing hippocampal amnesics can accurately identify trial-unique (never-before-seen) faces (3, 43-45), in contrast to limited evidence suggesting hippocampal amnesics were impaired at recognizing trial-unique kaleidoscopic images (46). Our data argue against the idea that the hippocampus only supports repeatedly-used stimuli, as a large proportion of neurons differentiated novel from repeat videos throughout the entirety of recording sessions despite the use of never-before-seen videos on each trial. We note that videos used in a given session were drawn from the same >40, 2-3 minute source videos, making it possible some content was similar across trials, although effort was made to avoid similarities by selecting for videos with many cut scenes like music videos and movie trailers and also making sure 5 s on either side of every 2 s snippet used in the task was not used at any other point. In addition, our previous work also found a large proportion of monkey hippocampal neurons that clearly distinguished novel from repeat images despite images being carefully selected to be trial-unique (23). Since in that work lure (similar) images were used in addition to novel and repeat images, and the task was not technically free-viewing due to images disappearing when monkeys looked away from the image (see also (22)), it was possible that hippocampal involvement might have been due to the pattern separation (47) or eye control (28) aspects of the task, respectively. Now, considering our results in addition to these previous studies (22, 23), the most parsimonious explanation is that the hippocampus supports the familiarity component of recognition memory for previously unseen stimuli.

The majority of recognition memory studies recording single hippocampal neurons in monkeys have required explicit identification of repeated stimuli—typically via a touch (18, 19, 21) or lick (20). These studies found few (<5% (20)) or no (1) hippocampal neurons significantly responded to repeated images. However, recent studies (22, 23) using preferential looking paradigms to detect memory—where monkeys were serially presented images and only trained to remove their eye gaze from the image to cease presentation—have shown large proportions (∼50%) of hippocampal neurons significantly change in firing rate to repeat images (summarized in Table 1). Hippocampal recordings had not been performed while monkeys watch relatively natural video stimuli, and therefore it is unknown if the recognition memory signals during these tasks were due to the lack of an explicit response component or related to the volitional control of eye gaze, which has been frequently linked to hippocampal activity (28). Here, we find a significant proportion of hippocampal neurons that show recognition memory responses to freely-viewed, fixed-duration videos, with the detection of repeat v. novel videos time-locked to the duration of each of the key conditions: subclip 1 and subclip 2 (Fig. 3). Therefore, the engagement of hippocampus is most likely due to the free-viewing nature of our task with no explicit memory probe linked to reward. Notably, work measuring looking times in a preferential looking framework with human hippocampal amnesics found their memory for repeat images impaired in comparison to controls (48), supporting the importance of the hippocampus in free-viewing recognition memory.

While the hippocampus has not been shown to support recognition memory in monkeys work utilizing explicit response tasks, a handful of electrophysiology studies in human patients do show evidence of recognition memory in absence of recollection. One pair of studies measuring single hippocampal units found evidence of recognition memory via similar numbers of neurons with both repetition suppression and repetition enhancement responses even when human patients failed at recollection (10, 42). However, the task design in these studies involved an object-in-place (49) learning paradigm, which is likely to engage the hippocampus via its known role in associative learning (36, 49), making it difficult to directly compare with our free-viewing video task. Another study measured iEEG high-frequency activity as a proxy for neural firing rates, and found the hippocampus contributes to both recollection and familiarity to correctly-identified, repeated words (9). However, the recognition memory judgments were only done after free recall periods for each list of words, likely inducing recollection for many of the words that were later recognized. One other study had humans identify repeated images in serially presented blocks of faces or scenes, and found a large proportion of single hippocampal units that responded to previously seen stimuli, including 31% via repetition enhancement (11). The authors argue that the disparity between their study and past monkey work might be due to recollection of previously seen or familiar items (11), but as in the other human studies are likely seeing some combination of recollection with a weaker recognition signal for recently-seen stimuli(50). Overall, despite the task differences, neural evidence for recognition memory in human hippocampus is likely to be related to the responses shown in our task.

Surprisingly, unlike in the studies mentioned above, the majority of video-responsive neurons showed repetition enhancement to repeat videos (∼70% of neurons to either subclip 1 or subclip 2, Table 1). This proportion is also unlike previous serial image studies in monkeys, where only a minority of hippocampal neurons showed repetition enhancement (∼40% of neurons in both studies, Table 1). A minority of repetition enhancement neurons is also typical during DMS tasks in other medial temporal lobe regions like inferotemporal/perirhinal (51, 52) and entorhinal cortex (53). We suggest that the use of video stimuli in our task might be responsible for this previously unseen pattern of responses. Videos contain series of object, faces, and scenes within a context, and using such life-like stimuli that have frequently driven hippocampal activity in human studies (33-35, 54) are likely to yield neural correlates indicative of the naturalistic experience of everyday life (55). In addition, videos introduce an element of time, and hippocampal neurons have shown strong evidence for roles in elapsed time (56), temporal order (57), and associative memory with a timing condition (58). To understand the relationship between hippocampal repetition enhancement and familiarity, future work would ideally design a task to parametrically assess the degree to which this pattern of single neuron responses reflects behavior, as has been done for repetition suppression in inferotemporal cortex (16).

To the best of our knowledge video stimuli have not been used in electrophysiological studies of monkey memory. However, videos have been used in a human electrophysiological (iEEG), single-unit study of memory (33) where experimenters selected for hippocampal or entorhinal cortex neurons responsive to long-term memory “concepts”. Videos of people (e.g. Tom Cruise) or content (e.g.

The Simpsons) meaningful to the patients were screened and these single neurons selectively rose in firing rate while viewing or later recalling those specific videos. In our results, the hippocampal neurons respond to a large proportion of repeated videos seemingly agnostic to the content of each video, as the change in firing rate for these neurons is clear after averaging across trials despite the use of unique videos with variable content on every trial (Fig. 4). It is unlikely the repetition enhancement we see reflects single neurons selective for many concepts, even if monkeys may have concept cells (59), as only 5.7% of videos elicited sustained responses in single hippocampal or entorhinal units in the human video task (33).

Further, we do not consider the hippocampal responses we see to be related to episodic memory, which we expect to be a sparse code(60). Instead, the responses seem more likely related to a familiarity signal for freely-viewed stimuli.

The responses during the two novel v. repeat task conditions in each retrieval period further confirm the general nature of this familiarity signal. The classifier trained on hippocampal population responses in one subclip period could decode which trials shown to the monkeys were repeats in the other period (Fig. 3), even though the videos shown in subclip 1 and subclip 2 were different (deliberately, videos from the first and last two seconds of the clip were never from the same source videos). Adding to the generalness of this recognition signal, the neurons responsible for classification via repetition enhancement in one subclip period tended to be the same ones responsible for classification in the other (Fig. 5C). In addition, despite both subclip 1 and subclip 2 showing the same pattern of responses to repeat videos, this signal did not hold across the delay period (Fig. 2D, 3). This absence of persistent firing suggests the repetition enhancement signal is not related to hippocampal working memory as has been shown in numerous explicit tasks (61-64). It remains possible that working memories are still present in dynamic or activity-silent states that our classifier would not detect, however(65).

Finally, despite designing the task to potentially cue recall during the delay by showing repeated videos in subclip 1, we found no evidence for recall during the delay period (SI Appendix, Supplementary Text & Figs. S1-S2). Since concept cells, working memory, and recall are all unlikely to explain the pattern of responses we see to repeated videos, the simplest explanation is these single hippocampal neurons support familiarity for recently-seen videos during free-viewing.

The boundary conditions for hippocampal activity during recognition memory have long been unclear, as findings in single neuron monkey work (1) and human amnesic research often seem contradictory (3, 6). Our work helps to resolve some of these contradictions. Evidence of hippocampal activity during recognition of unfamiliar (trial-unique) videos argues against the idea that the hippocampus is recruited only for recognition of reused stimuli (3). Instead, our results are in line with a number of studies of hippocampal amnesics with recognition memory impairments for most stimulus types other than faces (3, 44-46). We also suggest that the use of a free-viewing task is important for the recruitment of the hippocampus in the familiarity aspect of recognition memory. Most recognition memory research (1)—especially in humans (3)—requires retrieval probes to explicitly test participants’ memory, while studying hippocampus during free-viewing of naturalistic videos may have been overlooked due to the task’s seeming simplicity. We believe such naturalistic paradigms may be key to further understand how and when the hippocampus contributes to human memory.

## Methods

### Animals

Two female rhesus macaque (I = 6.1 kg, age 7; W = 6.5 kg, age 11) monkeys were surgically implanted with titanium headposts using aseptic techniques under general anesthesia and trained to be comfortable visually interacting with a computer monitor for juice reward while fixed in a primate chair for multiple hour recording sessions. All procedures and treatments were done in accordance with NIH guidelines and were approved by the New York University Animal Welfare Committee.

### Task

The task involved presentation of a series of videos with fixation “games” that rewarded the monkey before each video was shown. The game was to center fixate on a purple dot (radius = 1°, fixation maintenance required within 1.3° of dot center for 500 ms) to earn drops of 50% juice reward. Reward was never given during video presentation in order to separate reward from task-relevant stimuli. Monkeys were only required to fixate their eyes during these fixation games and during pre-video periods, which occurred before the start of videos to ensure the monkeys’ attention was drawn to the center of each video before it was presented. Pre-video dot properties were identical to fixation game dots except the color was changed to pink. Once pre-video fixation was achieved a video was shown (either clips 1-3 or subclip 1), and monkeys were permitted to freely view them, thereby allowing us to use natural viewing as a gauge of their interest in a given video (preferential looking).

The encoding phase of each trial began with a fixation game, where monkeys earned 0.35mL of 50% juice for achieving fixation. Next, a pre-video dot appeared, and once the monkey achieved fixation on this dot for 500 ms, clip 1 was shown. Then, after a second fixation game and pre-video fixation, the same 6 s video was repeated as clip 2. Finally, after a third fixation game and pre-video fixation, the same 6 s video was repeated as clip 3. A blank screen was shown for 450 ms after clips 1-2 before the dot reappeared for the next fixation game. A blank screen was shown for 300 ms after clip 3 before entering the retrieval period.

Clips were comprised of 3, 2 s videos shown contiguously as a single, 6 s episode (19.1° x 14.3° [640 x 480 pixels]). We refer to the 3 contiguous videos that comprise each clip as a, b, and c. Clips 1, 2, and 3 were identical videos meant to improve encoding through repetition. The videos were drawn from a new set of >40, 1-3 min source videos each daily session and presented without sound. The source videos were taken from Internet videos (YouTube) of scenes involving animals, people, animations, and other such moving subject matter. Music videos with a changing narrative (not focused on the same scene) and trailers to movies were typically used due to their short length and fast-moving, non-repetitive content. A custom Presentation (Neurobehavioral Systems) program was written that would randomly select 2 s snippets from that session’s source videos, only take snippets at least a 5 s buffer from any previously seen snippet, and ensure each snippet would never be used again. While there was a chance of similar clips across trials since we used the same source videos each session, the short scenes of the music videos/trailers as well as the use of the buffer minimized overlapping content. In addition, since each clip was always comprised of 2 s snippets from 3 different source videos, the clip itself was almost guaranteed to be a unique construction of content sources since even with only 40 source videos there are 40^3^=64,000 order arrangements, while monkeys averaged only 125.9±2.5 trials per session.

The retrieval period of each trial began with a fixation game and a pre-video period before subclip 1 was shown. Immediately after subclip 1 was shown, a larger (1.6°), teal dot was shown in the center of the screen for 1 s. The monkey was not required to fixate to this dot, which served merely as an indicator of the delay period between subclips. Finally, subclip 2 was shown followed by a 3 s inter-trial interval.

Subclips were comprised of 2 s videos surrounding the 1 s delay period. Three trial types with a different combination of subclips were shown at equal proportions: i) subclip 1 a repeat of the first 2 s of the clip (i.e., section a) and subclip 2 a repeat of the last 2 s of the clip (i.e. section c), ii) subclip 1 a repeat of the first 2 s of the clip and subclip 2 a novel (unseen) video snippet from the source videos, and iii) subclip 1 a novel video and subclip 2 a novel video. The delay period was only set at 1 s—instead of 2 s to better match the structure of the clips—owing to pilot behavioral data indicating delays longer than 1 s made it more difficult for monkeys to maintain attention.

### Behavior

We used looking times at the videos as a proxy for memory of the repeated videos (preferential looking). Monkey eye scan paths were recorded using an infrared camera (IScan, Inc.) as the animals viewed a 19-inch LCD monitor (Dell) positioned 0.55 m from the left eye to the center of the screen. Monkeys were considered to be looking at the video when their eyes were within the bounds of the 640×480 pixel video plus a 20 pixel buffer surrounding the perimeter of the video. We then measured the proportion of the time that the monkey looked within the bounds of the clip or subclip videos to find looking times. We then set a threshold to define preferential looking, where if the monkeys looked longer at novel subclips than clip*threshold, we considered them to have identified the novel video (true positive). And if the monkeys looked shorter at repeat subclips than clip*threshold, we considered them to have identified the video as a repeat (true negative).

To give a concrete example, if the monkey looked within the bounds of clip 1a for 1.6 s, and the monkey looked within the bounds of a novel subclip for 1.4 s, this subclip would be a true positive, since 1.6*0.8=1.28 s is less than 1.4 s. But if the monkey looked at the novel subclip for 1.0 s, the video would be considered a false negative (1.0<1.28 s).

Trials were only considered for analysis if the clip period was looked at for at least 0.5 s of the 2 s segment. d’ values were calculated using the formula d’ = norminv(true positive rate)-norminv(false positive rate), where norminv is the inverse of the normal cumulative distribution with mean of 0 and standard deviation of 1. We set a constant threshold of 0.8, which we found worked well for both key conditions outlined below, although thresholds in the 0.6:0.9 range all gave results within 10% for both subclip 1 and subclip 2 d’ values.

We performed two comparisons to analyze behavior: clip 1a (0-2 s) v. subclip 1, and clip 1c (4-6 s) v. subclip 2, since these were the two segments of clip 1 that videos could be repeated in the retrieval period. The same comparisons were made using clip 3 instead of clip 1 as well with the same design. Statistics for comparisons of average looking times (Fig. 1B) for each monkey were Bonferroni-corrected for family-wise error rate (FWE) by adjusting the p-values for the 6 total comparisons (6*p-value). Since the p-value for monkey I was significant when testing clip 1a v. novel videos in subclip 1 but monkey W’s was not, we ran a mixed effects regression to test if Monkey I’s preference for novel subclip 1 videos was significantly greater than for Monkey W. This regression was of the form: Looking time ∼ monkey + period + monkey*period, with the p-value for the interaction monkey*period the key value to test this hypothesis.

### Electrophysiology

Once monkeys were trained on the task, MRI-designed, custom-fit circular HDPE plastic recording chambers (Rogue Research) were cemented to their skulls with acrylic perpendicular to the horizontal plane in stereotaxic coordinates and dorsal to their right hippocampi (chamber outlines are shown in SI Appendix, Fig. S4). Single-unit recordings were performed by lowering glass-insulated electrodes (∼1-MΩ impedance; Alpha Omega) or glass-insulated tetrodes (∼0.5-MΩ impedance for all four shafts; Thomas Recording) via a hydraulic (Kopf Instruments) microdrive through 23-gauge metal guide tubes that reached from the chamber grid to ∼10–15 mm dorsal to hippocampus. Guide tubes were placed at new locations at the beginning of each week and removed at the end of the week after multiple sessions of recording. Neural signals were acquired, filtered, amplified 1,000 times, and digitized via the MAP Data Acquisition System (Plexon) at 40,000 kHz resolution. Single unit waveforms were sorted manually using a combination of offline sorter (Plexon) and automatic clustering [Wave_Clus (66)], with all clusters from the latter checked manually in offline sorter.

### Single unit analysis

We recorded a total of 209 neurons across the 116 sessions detailed in our behavioral analysis. All of these neurons were used in our population sparseness analysis, which was performed as in previous work (23). Briefly, population sparseness is a measure of the proportion of neurons in a population that respond to a given stimulus (37). Ideally, this measure is done for single stimuli repeated multiple times across trials, but since by design we do not repeat stimuli across trials, we measured the sparseness across videos shown during subclip 1 for all trials (therefore, instead of pooling trials for the same stimulus, we pooled trials for the same task period). The equation is:

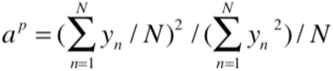

where yn is the average firing rate for each neuron and N is the total number of neurons. The population sparseness value of *a*^P^ = 0.30 was similar but slightly more sparse than values for monkey hippocampal neurons firing to repeated image stimuli (*a*^P^ = 0.33) in previous work (37).

To perform the single unit analyses in Tables 1-2, we kept only those neurons with a minimum average firing rate of at least 0.5 Hz across all trials. The number of neurons used in each analysis are listed in the table, with neuron inclusion based on a minimum of 25 trials for all tests involving subclip 1 (yielding 144 neurons) and 20 trials for all tests involving subclip 2 (yielding 156 neurons). That is, there were at least 25 novel and 25 repeat trials recorded for each neuron during the subclip 1 period, and at least 20 novel and 20 repeat trials recorded for each neuron during the subclip 2 period (even though novel and repeat trials were evenly distributed for each subclip, trial numbers for each condition for each neuron are variable due to monkeys occasionally stopping work early or neuron isolation being lost while the task was running). Spikes in the window of 0.1-2.1 s were measured separately for each condition (novel and repeat). Neurons with significant increases or decreases in firing rates (Table 1) were found by performing a Wilcoxon rank-sum test (p<0.05) between trials from a given condition and firing rates pooled across all fixation games (there are 4 such fixation periods per trial—before each clip and subclip 1). Neurons with significant differences between conditions (Table 2) were found using the same test. Chi-square tests were used in both tables to assess the likelihood of one group of neurons being larger than another (e.g. increase v. decrease in firing rate to repeats), with p-values being Bonferroni-corrected to account for family-wise errors (FWE) by the number of tests within each table (i.e., 4 tests for statistics in Table 1, and 2 tests for statistics in table 2). For the comparison of novel v. repeat trials using behavior, we found the repeat videos identified as repeats and the novel videos identified as novel from our behavioral analysis above and then ran the same tests as in Table 2. We reduced the trial minima to 15 (yielding 152 neurons) and 10 (yielding 147 neurons) trials for subclip 1 and 2, respectively, and used the same FWR correction of the chi-square p-values for 2 tests.

**Table 2.**
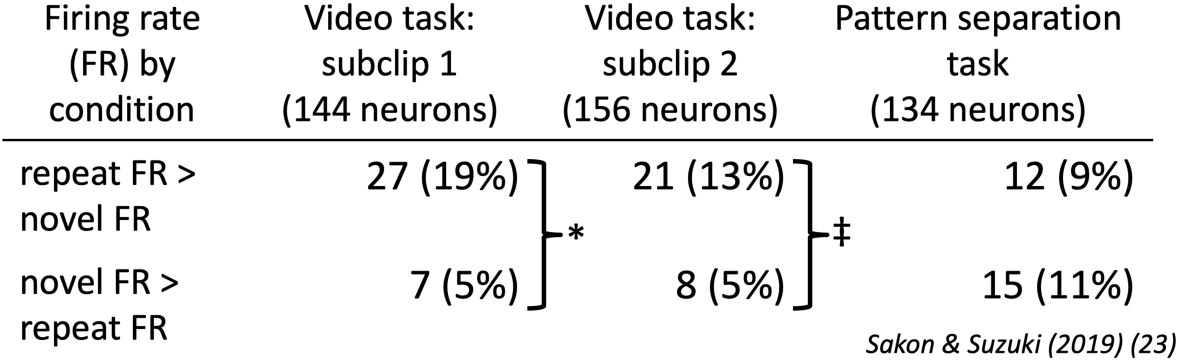
Comparison of novel v. repeat responsive neurons between video task and image recognition task. The number (%) of monkey hippocampal neurons (with FR≥0.5 Hz during given period) that respond with significantly stronger firing rates (p<0.05, Wilcoxon rank-sum) between novel and repeat stimuli for given for each task. *p=1.0e-3, ‡p=0.04, FWE-corrected.

### Population classifier analysis

We performed a 10-fold, cross-validated, L2-regularized (ridge) logistic regression for the neural firing rates in our population with fixation period (baseline) firing rates >0.5 Hz. This setup means the classifier was trained on 90% of trials and tested on the remaining 10%, and this process was repeated 10x in a balanced design such that every trial was part of the test set once.

Predictors were the identity of the subclip 1 video (novel or repeat) with the response variable the firing rates for each neuron. Separate classifiers were trained on each time bin, with 500 ms spike integration windows and 250 ms step sizes for Fig. 2A-C and 200 ms integration windows and 100 ms step sizes for Fig. 2D. We used regularization since with small numbers of trials on each run perfect classification on any given training or test set would weight that neuron to infinity. Ridge regression was used since our goal was to find all neurons that contributed to classification and not remove redundant signals. By keeping weights for each neuron, we could also gain an approximate rank order of each neuron’s contribution to the classifier (Fig. 5). The regularization parameter (λ) was set to 0.15, although a range of numbers (from 0.05 to 0.3) yielded similar results.

We attempted to use a similar number of neurons in each classifier we trained, with the number of neurons differing depending on the minimum number of novel and repeat trials we required for each analysis. For the classifier of subclip 1 identity in Fig. 2A (and also 2D), where we used all trials, we required at least 40 novel and 40 repeat trials for each neuron, which gave a population of 96 neurons. For the classifier of subclip 1 and subclip 2 identity in Fig. 2B & Fig. 3, where we eliminated the (1/3^rd^ of) trials which were repeat then novel for subclips 1 and 2, we required at least 35 novel and 35 repeat trials for each neuron, which gave a population of 86 neurons. Finally, when we combined trials across subclips 1 & 2, we required 100 novel and 100 repeat trials, which gave a population of 106 neurons.

The prediction accuracy for each classifier is an average over many permutations sampled from different trials in the same group of neurons. Since the response variable for the classifier trained at each time bin is a neuron X trial matrix (e.g., for Fig. 2A, 96 x 80), and many of those neurons will have more than the minimum number of trials we set for that classifier, we randomly subsampled trials for those neurons with more than the minimum. We ran the classifiers in Fig. 2A, 2B, and 2D for 200 permutations, averaged across those permutations to find the prediction accuracies at each time bin, and used the 5^th^ and 95^th^ percentile classification runs for the error bars shown in Fig. 2. For computational reasons due to the larger number of trials, we only used 80 permutations for the classifier in Fig. 2C. Notably, the prediction accuracies were remarkably consistent across multiple runs for all four classifiers.

To find periods of significant classification, we used a cluster-based metric that corrects for multiple comparisons across time bins (39). For each of the 200 (or 80) subsampled permutations described in the last paragraph, we ran 5 additional permutations of the classifier with the labels (novel and repeat) shuffled, thereby creating a surrogate distribution of 1000 (or 400) total permutations. A maximum cluster was found for each of these surrogate permutations, where a cluster is the sum of the prediction accuracies for a stretch of contiguous time bins that cross a threshold. The threshold can be set to any value without increasing the false alarm rate, but a threshold set too high or low will decrease the sensitivity (39). We used 0.625 as the threshold for the classifiers in Fig. 2A and 2C, and 0.6 for the classifiers in Fig. 2B and 2D. Next, we took the real prediction accuracies, from averaging across the 200 (or 80) permutations, and found all contiguous clusters above threshold as we did for the surrogate data (in this case, not just taking the maximum cluster). Finally, we found which of these cluster values in the real data were greater than the 95^th^ percentile cluster value in the surrogate data, and any such clusters above this value were considered significant. Significant clusters of classification in Fig. 2 are denoted with asterisks, with all asterisks in neighboring time bins indicating a single, contiguous cluster.

For the heatmap of prediction accuracy in Fig. 3, we trained and tested at each combination of time bins from 1000 ms before subclip 1 on until 3000 ms after subclip 2 off. The classifier at each point was identical to the classifier in Fig. 2B, with the diagonal of the heatmap being equivalent to the prediction accuracy in 2B. The prediction accuracies shown are the averages of 100 permutations with different subsamples of trials as described for Fig. 2B.

To find the neurons that most contributed to classification we see in the subclip 1 and subclip 2 periods (Fig. 4 and SI Appendix, Fig. S5), we ran separate classifiers on a single time bin from 500-2500 ms after subclip 1 on and from 500-2500 ms after subclip 2 on. These weights provide the ranks of which neurons had the most positive weights that contributed to the classifier. The distribution of weights for the classifier for the two subclip periods also fill the distributions used in Fig. 5.

### Cued recall analysis

We investigated neural responses during the delay between subclips with two analyses. First, we used a classifier in Fig. 2D similar to the classifier in Fig. 2B, but including all trials (the classifier in 2B removes the repeat/novel trial type for subclip 1/subclip 2). The predictor for this classifier was if the video in subclip 1 was a repeat or novel video, since we expect recall to only occur during the delay period after the monkey is cued with a repeat video—and not a novel video—in subclip 1.

Second, we looked at the trial-by-trial correlation (Spearman) of firing rates for each neuron when comparing the delay period with the portion of the clip we except to be recalled. That is, if a repeat video is shown during subclip 1 and this (4^th^ presentation of 0-2 s of that video after the clip was shown 3x) successfully cues recall in the monkey, we expect neurons during the delay period to recapitulate firing rates on those trials.

However, if a novel video is shown during subclip 1, that should not elicit recall, and we should not expect correlated firing between the clip and delay periods on those trials. For the delay period in these comparisons, we found the firing rate in the window from 0.1-1.1 s after subclip 1 off. For the clip period, we found the firing rate in the window from 2.1-4.1 s after clip on (since clip b should be what is being recalled after the monkey is cued with a repeat of the first 2 s of the clip, as diagrammed in Fig. 6A).

Statistical comparisons were all multi-level ANOVA, where we correlated out differences between monkeys and the interaction factor to isolate only the differences in variance between the correlations.

We measured 7 trial-by-trial correlations of firing rates this way and averaged them across neurons for Fig. 6B. For the first two comparisons, we correlated the middle 2 s of clip 1 with the delay period separately after a repeat video (post-repeat, when we expect recall) and after a novel video (post-novel, when we don’t expect recall). We did the same comparisons for clip 3 v. delay as well.

Next, we ran 3 controls to get an idea of the correlation values we should expect across our hippocampal population. First, a negative control where we compared firing rates to videos in unrelated periods. We found the firing rates from 0.1-2.1 s after presentation of clip 1 to firing rates during subclip 2, where a video is presented that is never a repeat of the first 2 s of the clip (since subclip 2 can only be a presentation of the last 2 s of the clip or a novel video). Next, we performed a positive control where we compared the same segment of video between two different clips. We used the firing rates from 0.1-2.1 s after video presentation during both the clip 1 and clip 3 periods.

Finally, we performed a positive control where we compared firing rates to different segments within the same clip, even though different videos were shown in these segments. For this we correlated the firing rates between 0.1-2.1 and 2.1-4.1 s after the presentation of clip 3.

## Acknowledgements

We thank Andrea Shang, Ellen Wang, and Joyce Ho for expert animal care and Yuji Naya for suggestions in task development. This research was supported by the National Institutes of Health (R01-MH084964) and the Simons Collaboration on the Global Brain (542997).

## Author Contributions

JJS and WAS designed the experiment and wrote the paper. JJS collected the data and designed and performed the analyses.

## Data availability

Full data, or data pre-processed for a specific analysis, is available upon request.

## Code availability

Analysis code (in Matlab) and pre-processed data to reproduce analyses and figures are available on request.

## Supplementary Information for

## Supplementary text

### Preferential looking at typical 1.0 threshold still classifies novel from repeat videos

Preferential looking is typically used as a proxy for memory(1), albeit typically with a threshold of 1.0, meaning only subclips with looking times less than novel clips would be classified as repeats. This is a stricter interpretation of novelty preference and is generally used for side-by-side comparisons(2), but as a control we ran the same analysis with 1.0 instead of 0.8 to ensure our methods are sound even in this stricter framework.

#### First, our data using a threshold of 80%

Clip 1a v. subclip 1 (average across sessions for all data, with SE for all errors): 85.0±0.9% hit rate (true positives, i.e. novel videos classified as novel) 66.4±0.8% false alarm rate (false positives, i.e. repeat videos classified as novel) d’=0.70±0.05

Clip 1c v. subclip 2:

80.2±0.9% hit rate

54.1±1.0% false alarm rate

d’=0.80±0.05

Clip 3a v. subclip 1:

85.2±0.9% hit rate

66.7±0.7% false alarm rate

d’=0.72±0.05

Clip 3c v. subclip 2:

85.5±0.8% hit rate

68.8±0.8% false alarm rate

d’=0.63±0.04

##### Next, our data using a threshold of 100%

Clip 1a v. subclip 1:

56.2±0.8% hit rate

42.8±0.9% false alarm rate

d’=0.35±0.03

Clip 1c v. subclip 2: 63.7±0.9% hit rate

38.1±0.9% false alarm rate

d’=0.68±0.04

Clip 3a v. subclip 1: 52.8±0.8% hit rate

41.8±0.8% false alarm rate

d’=0.28±0.03

Clip 3c v. subclip 2:

73.1±1.2% hit rate

52.8±0.9% false alarm rate

d’=0.59±0.04

As expected, our d’ values are not as high for a threshold of 100%, but they are still well above chance (note the small standard errors), and better than previous values we’ve seen classifying similar v. repeat images using looking times (d’=0.14)(3).

### Recall is not reflected by hippocampal neurons during the delay

We designed our task with the intent of creating a key delay period between the subclips to investigate recall. Our hypothesis was that after presentation of a repeat video during subclip 1, which would represent the 4^th^ presentation of the video shown in clip a (Fig. 1A), the monkeys would then recall the next segment of the clip (clip b). We hoped this cued recall paradigm would incite hippocampal activity during this video-free delay.

We explored three analyses to find evidence of recall during the delay: 1) a comparison of firing rates during clip b vs. the delay, where we expected this middle video of the clip to be recalled during the delay after the monkey was shown a repeat in subclip 1; 2) a comparison of firing rates between the delay period after a repeat was shown in subclip 1 vs. when a repeat was shown in subclip 2 (a prospective signal); 3) a comparison of firing rates between subclip 1 and subclip 2 for the trials with the best v. the trials with the worst behavioral evidence of memory.

For the first analysis, we took each of the same 110 neurons used in our classifier, and found a vector of firing rates during the delay period for each trial after a repeat was shown in subclip 1. Next, since we expect this 4th repeat of the same video to elicit recall of the remainder of the clip, we found a vector of firing rates for those same trials during clip 1b, and then correlated these vectors of firing rates for each neuron (Fig. S1A). For comparison, we correlated the clip 1b vector with the delay period on trials when subclip 1 was a novel video, in which case we do not expect recall of the remainder of the clip to be elicited. Our expectation is for higher correlations of clip 1b v. delay when a repeat video instead of a novel video is shown during subclip 1.

Contrary to our expectation for recall, the correlation between clip 1b and delay after presentation of repeat videos was no different than between clip 1b and delay after novel videos during subclip 1 (0.15 ± 0.02 v. 0.17 ± 0.02, Fig. S1B). We found the same trend when we correlated clip 3b to the delay period as well, with lower correlations after repeat videos compared to after novel videos (0.14 ± 0.02 v. 0.16 ± 0.02, Fig. S1B). These four vectors of correlations are not significantly different than if they were drawn from the same distribution (p=0.83, multilevel ANOVA). In fact, when we correlate clip 1a with subclip 2— two periods that never show the same video and therefore should not correlate—the value is similar to these other comparisons (0.13 ± 0.02, Negative control in Fig. S1B), indicating the delay period shows no evidence of recall beyond basal levels of correlated firing.

For the second analysis of recall, we considered if repeat videos shown during subclip 1 elicit something like a prospective signal (4) for subclip 2. In other words, after the repeat cue in subclip 1, hippocampal neurons might show evidence of recalling the repeat video that might be shown in subclip 2(clip c) during the delay. To test this, we took each neuron and measured the correlation in firing rates across trials between the delay period after subclip 1 was a repeat v. subclip 2. If a prospective recall signal for subclip 2 exists, the correlation should be stronger when subclip 2 was then a repeat than when subclip 2 was a novel video. Instead, these two correlations were not significantly different across neurons (0.24 ± 0.02 v. 0.19 ± 0.02, p=0.10).

The lack of recall signal in these two analyses could be due to a number of factors. First, trial-by-trial correlations of firing rates might be a poor metric for comparing the state of the hippocampal population. To test this idea, we can correlate the firing rates of the same clip across different periods (clip 1a v. clip 3a; positive control between clips in Fig. S1B) or the firing rates at different times within the same clip (clip 1a v. clip 1c; positive control within clips in Fig. S1B). The correlations within hippocampal neurons for these two positive controls were higher than the other four distributions, and were not expected by chance when compared to the other distributions (p=0.017, multilevel ANOVA). Therefore, the hippocampal population does show trial-by-trial evidence of returning to a similar state, but only during the same encoding video (clip 1a v. clip 3a) or within the same temporal context (clip 1a v. clip 1c).

For the third analysis of recall, we asked if classifying trials via looking times (Fig. 1B) could select for those subclips that were best-remembered and therefore most likely to show evidence of recall (5). We looked at only the 1/3^rd^ of trials when a repeat video was shown in both subclip 1 and subclip 2. Our hypothesis is that if the monkey recalled the video when shown a repeat in subclip 1, they would then show lower looking times when a repeat was shown in subclip 2. Therefore, we found the ratio between looking times for subclip 1 and subclip 2 across trials for each session, ranked them, and split them into terciles. The highest tercile being the best memory (biggest decrease in looking time from 1 to 2) and the lowest tercile being the worst memory. Finally, we found the trial-by-trial correlation in firing rates during subclip 1 and subclip 2 for each neuron for both the best and and worst terciles.

Our expectation is that if the monkey remembers the video in subclip 1, they will then recall the remainder of the video, therefore showing the same recall signal in subclip 2 and correlated firing rates between the periods. And when the monkey does not show evidence of recalling subclip 1, we should then see lower correlations between subclip 1 and subclip 2 firing rates. We found no significant difference in the subclip 1 and subclip 2 correlations between the best v. worst memory terciles (p=0.37, Wilcoxon rank-sum), with the average correlations for the worst actually higher than for the best (best tercile, 0.19, worst tercile, 0.23).

### Repetition enhancement neurons are not an artifact of baseline firing rates

For the analysis shown in Table 1, we use baseline-corrected firing rates, where the baseline firing rate is the average firing rate for the fixation periods that come prior to each clip and subclip 1 presentation (four per trial, Fig. 1). It is possible that the reason we appear to see a large proportion of repetition enhancement neurons is that during clip 1, when a novel video is presented, the firing rates might jump high above baseline, thereby making it possible for subsequent subclip firing rates to only appear high compared to baseline but instead just be differing levels of repetition suppression. To test this hypothesis, we plotted the z-scored firing rate for each neuron during clip 1, where the mean and standard deviation of the fixation period firing rates are used for the z-score (Fig. S3). Therefore, z-scores below 0 indicate lower firing rates during clip 1 compared to fixation while z-scores above 0 indicate higher firing rates during clip 1 compared to fixation. As shown, the neurons mostly showed lower firing rates during clip 1 compared to fixation, meaning neurons generally fire lower to novel video presentations at clip 1 than at fixation.

What about the key neurons that show the repetition enhancement effect? Of the 28 neurons in Fig. 5C that rise for both subclip 1 and subclip 2, only 7 of them show higher firing rates to clip 1 than fixation. Therefore, we have ruled out the possibility that the repetition enhancement effect we see during subclips 1 and 2 is actually lower in firing rate compared to novel video presentation at clip 1.

**Fig. S1.**
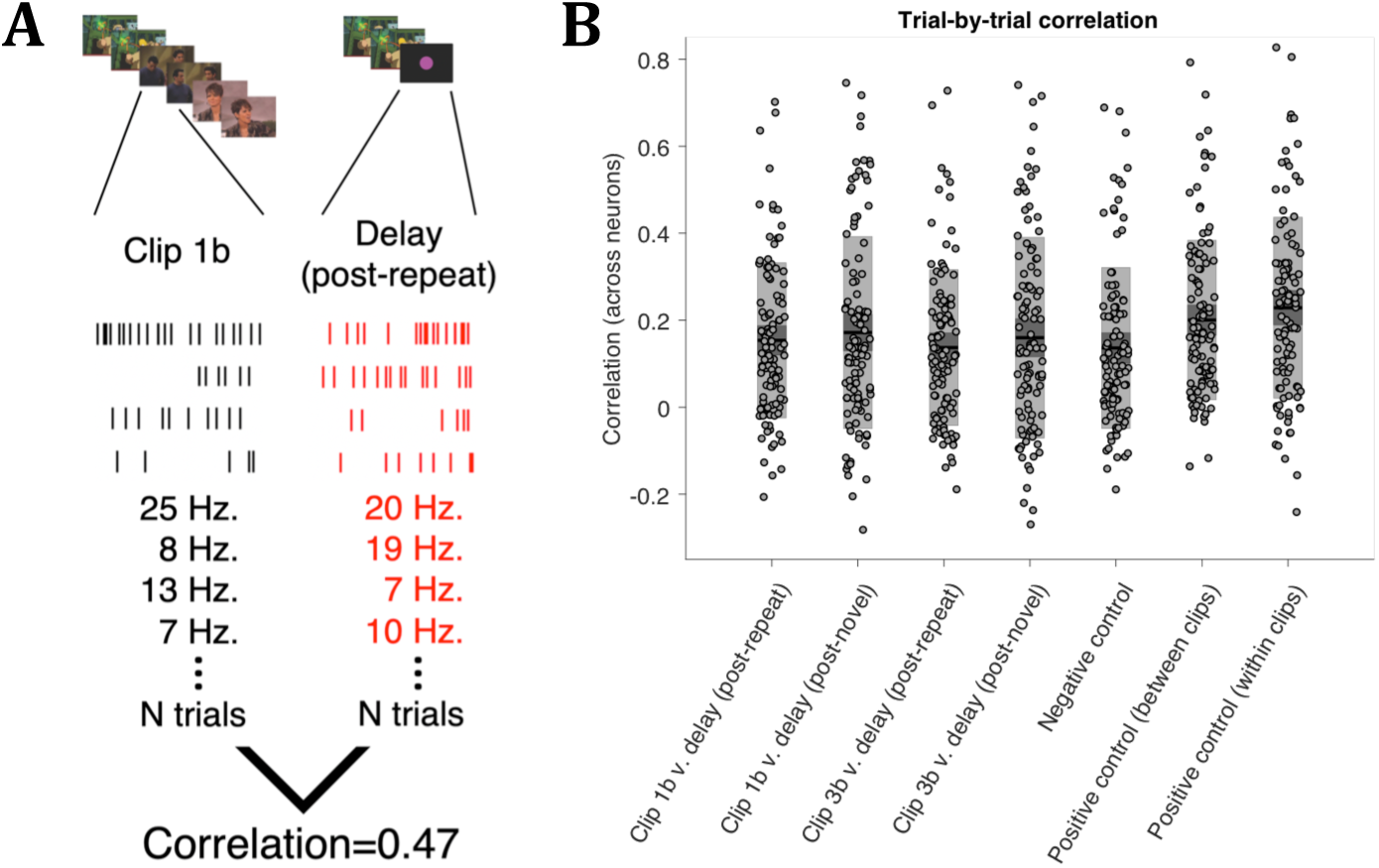
Hippocampal neurons show trial-by-trial correlations in firing rates within and between the clip periods but not between the clips and delay. **A)** Example correlation (Spearman) analysis for a single neuron. To test cued recall, vectors of firing rates during key periods for each neuron were correlated to find trial-by-trial evidence of the hippocampal population returning to a similar state. The expectation is that if the monkey is cued to recall the clip video, delay period firing rates after a repeat video is shown in subclip 1—but not after a novel video is shown in subclip 1—should correlate with the firing rates to the middle video of the clip. **B)** Average correlations across 110 neurons. The negative control is the correlation between unrelated videos, while the positive controls are the correlations between repeated clips or within the same clip. Shaded boxes are the 95% confidence intervals (dark gray) and the standard deviations (light gray).

**Fig. S2.**
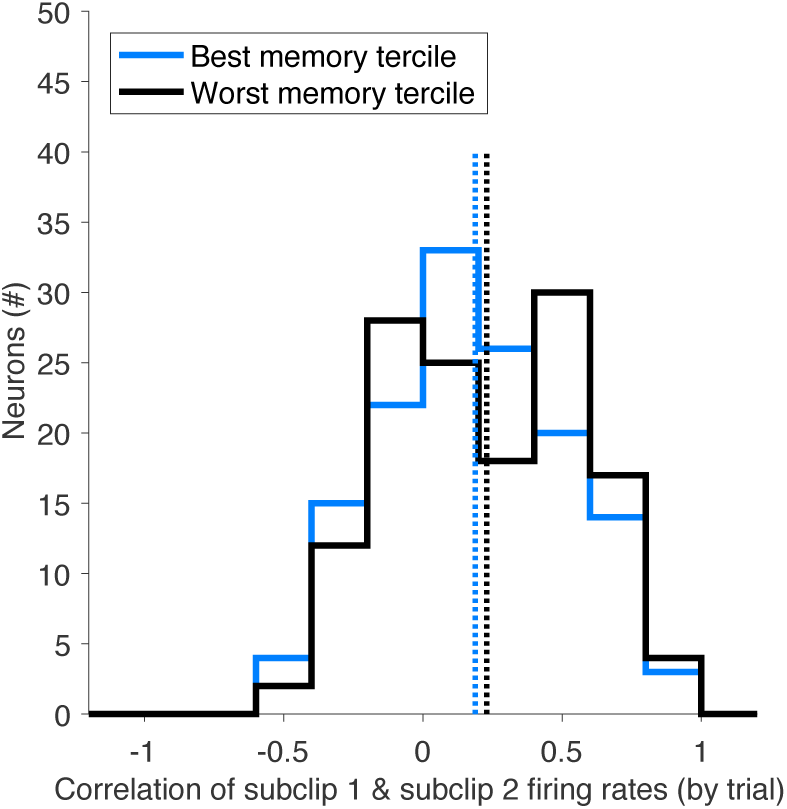
Trial-by-trial correlations in firing rates between repeat presentations in the subclip 1 and subclip 2 period are not different between the best and worst memory trials. We took each trial where a repeat video was shown during both subclip 1 and subclip 2 and found the best and worst tercile trials of memory for the second subclip based on looking times (a larger drop in looking time from subclip 1 to 2 indicates better memory). There was no difference between the distributions (p=0.37, Wilcoxon rank-sum).

**Fig. S3.**
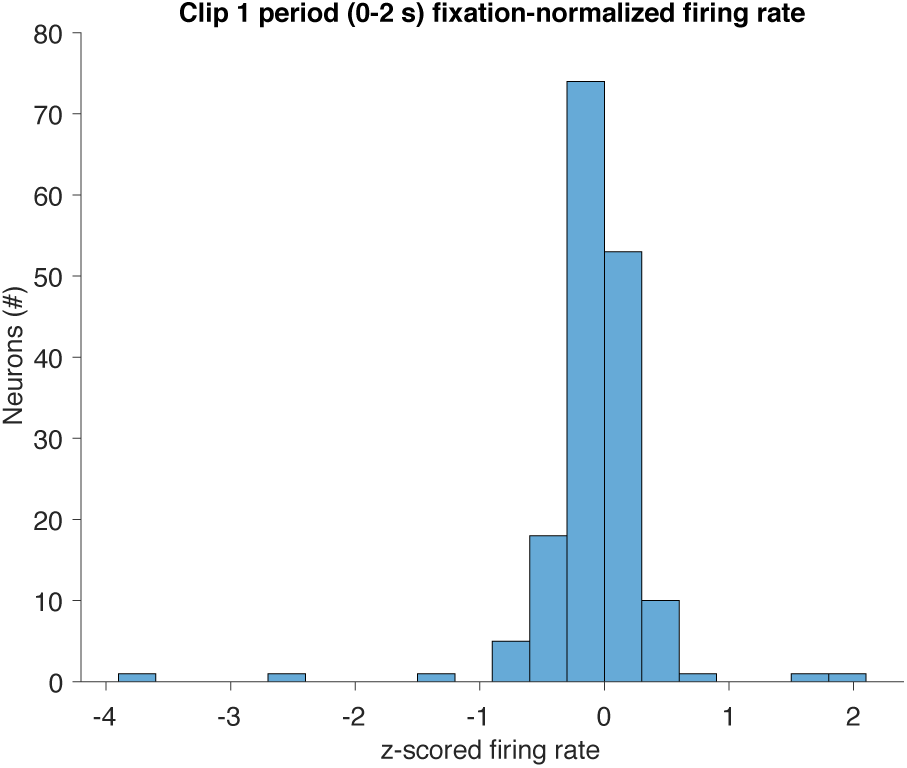
The majority of neurons show lower firing rates during clip 1 than baseline. For each neuron, we z-scored the firing rates from 0-2 s after video presentation in clip 1a (after including a 0.1 s offset) using the mean and standard deviation from the pooled firing rates from fixation periods on each trial. Z-scores lower than 0 indicate lower firing rates to clip 1 than baseline, while z-scores higher than 0 indicate higher firing rates to clip 1 than baseline.

**Fig. S4.**
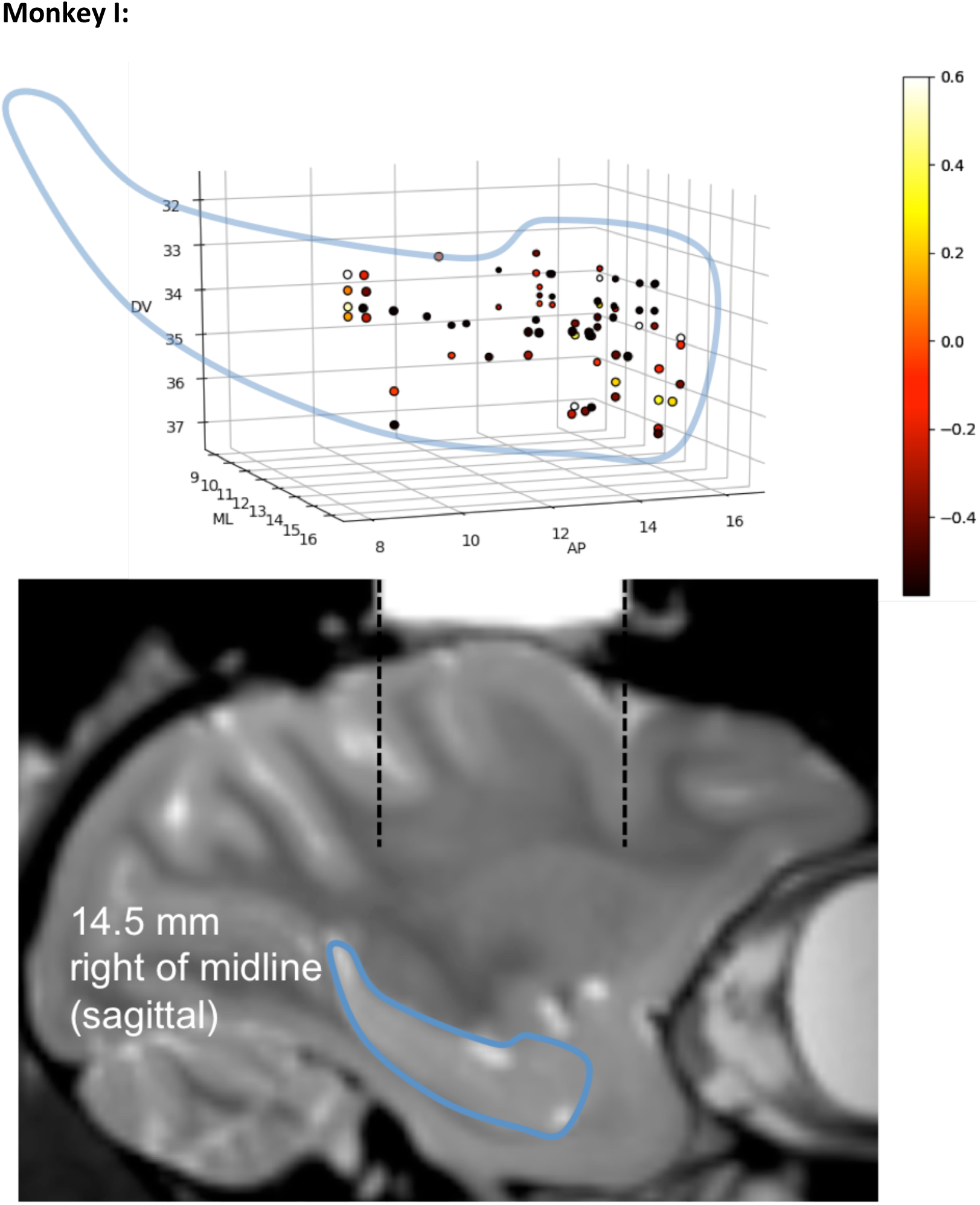
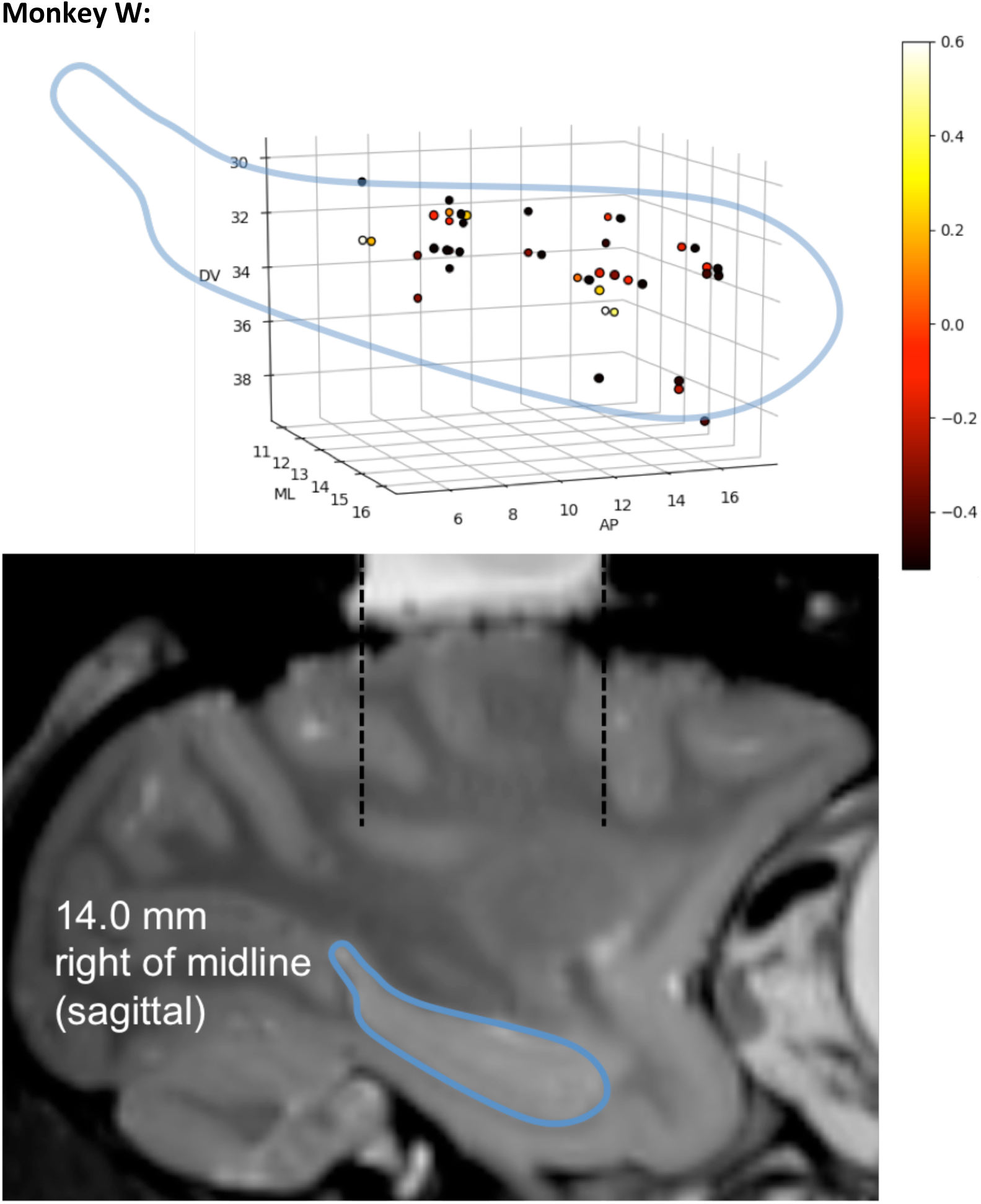
Hippocampal recording locations for each animal. Each point represents the recording site of a neuron. Top: The outline of the hippocampus, reproduced from the MRI image at bottom, gives an estimate of the relative locations within the hippocampus of each animal. The colors of each point are the weights from the classifier for Subclip 1 (Fig. 5A), and the sizes indicate the ML position with smallest being closest to midline (min=9.5, max=16). Note the heterogeneity of the weights across the hippocampus, with neurons showing strong repeat v. novel classification throughout the anterior-posterior extent in both monkeys. The top-6 weights (5 from monkey I, 1 from monkey W) were each set to 0.6 to increase the dynamic range of the colorscale. Bottom: Sagittal MRI images at indicated distance from midline with hippocampus outlined in blue. Dashed lines show the extent of the recording chambers. All electrodes were inserted through gridholes parallel to these lines. AP=anteroposterior. ML=mediolateral. DV=dorsoventral.

**Fig. S5.**
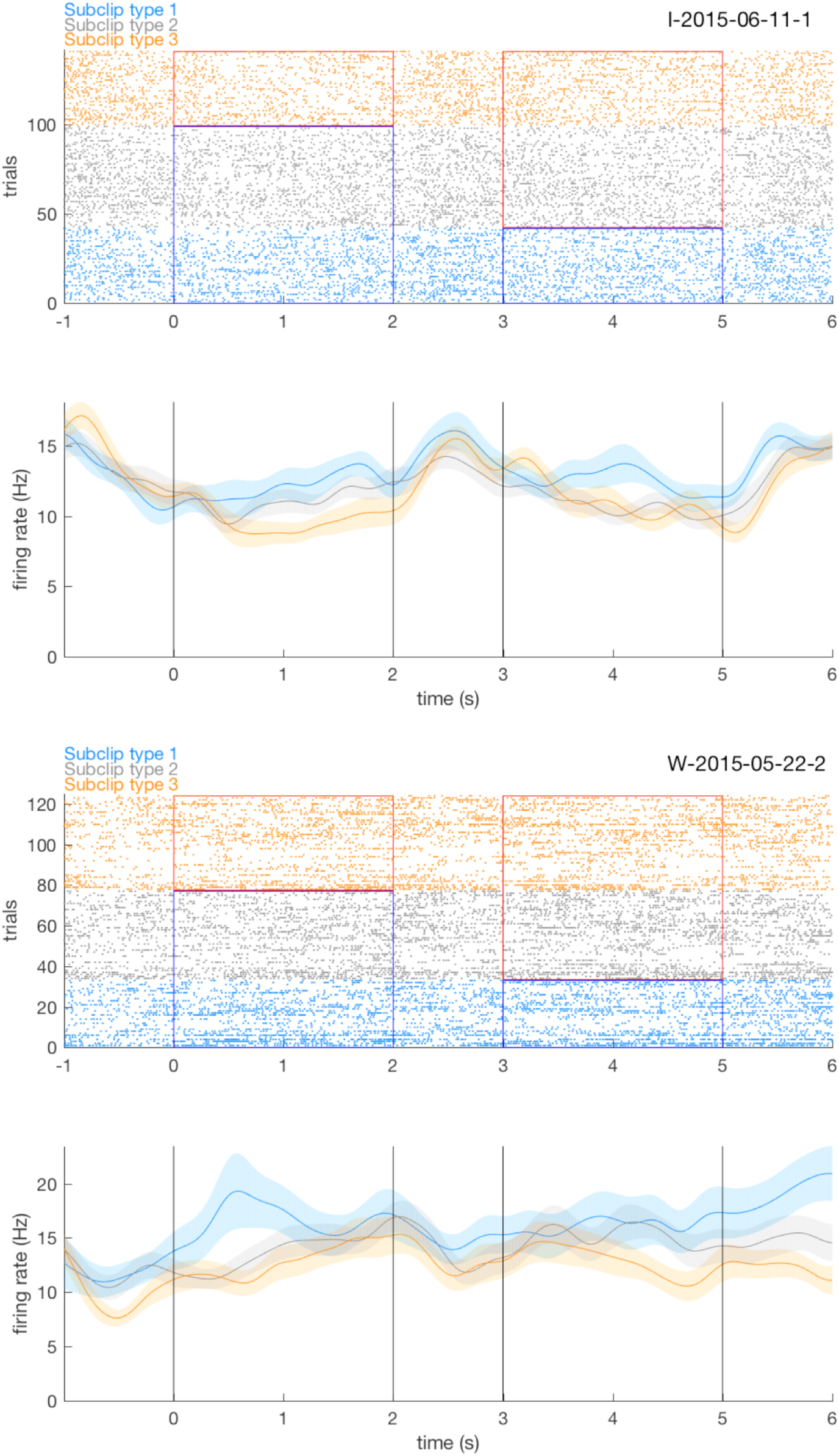
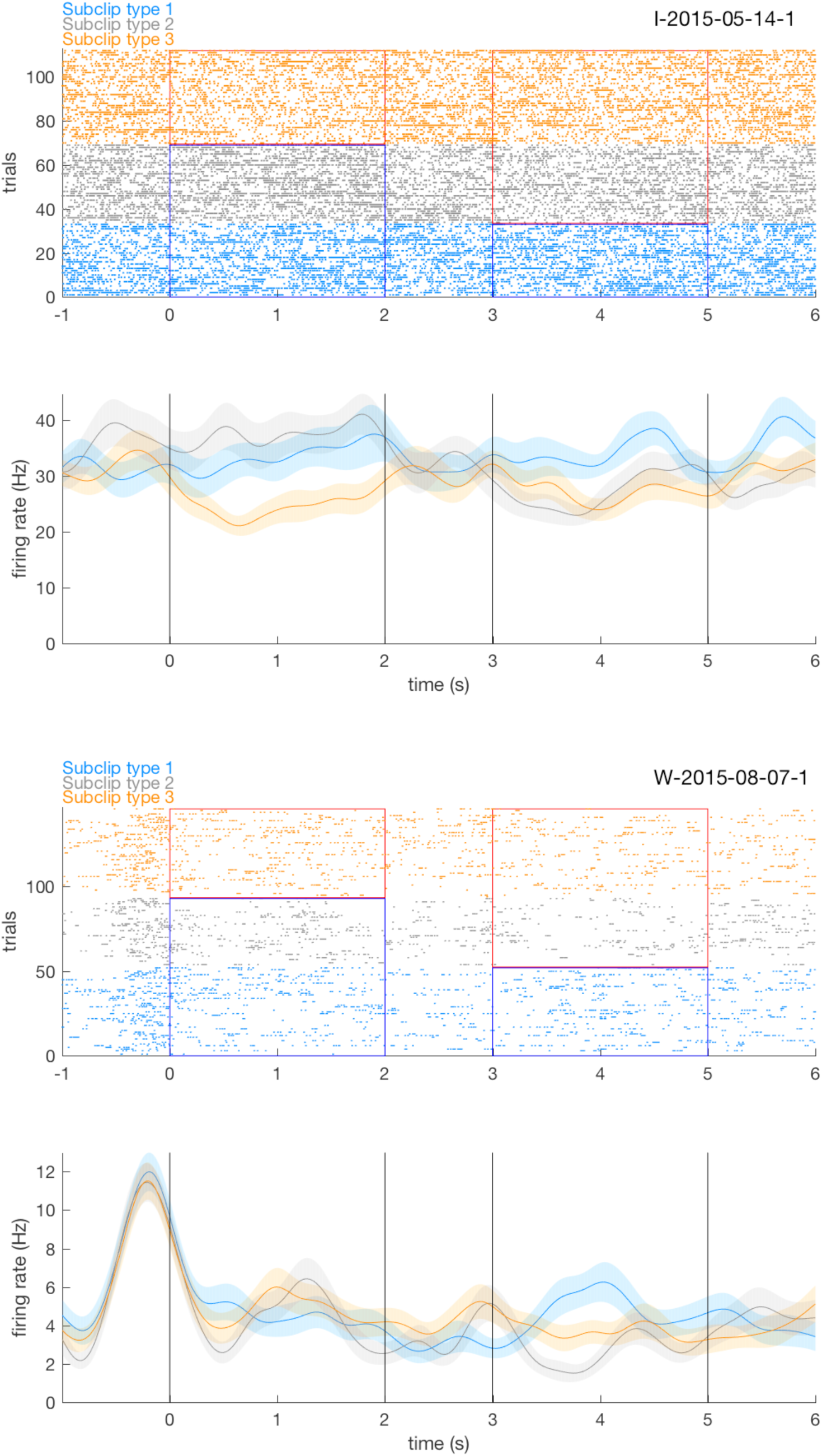
Additional examples of individual neurons. Conventions as in Fig. 4. These neurons were the 8^th^, 6^th^, 15^th^, and 22^nd^ most positive weights in the classifier for the subclip 1 period and the 9^th^, 22^nd^, 4^th^, and 14^th^ most positive weights in the classifier for the subclip 2 period (Fig. 2).

**Fig. S6.**
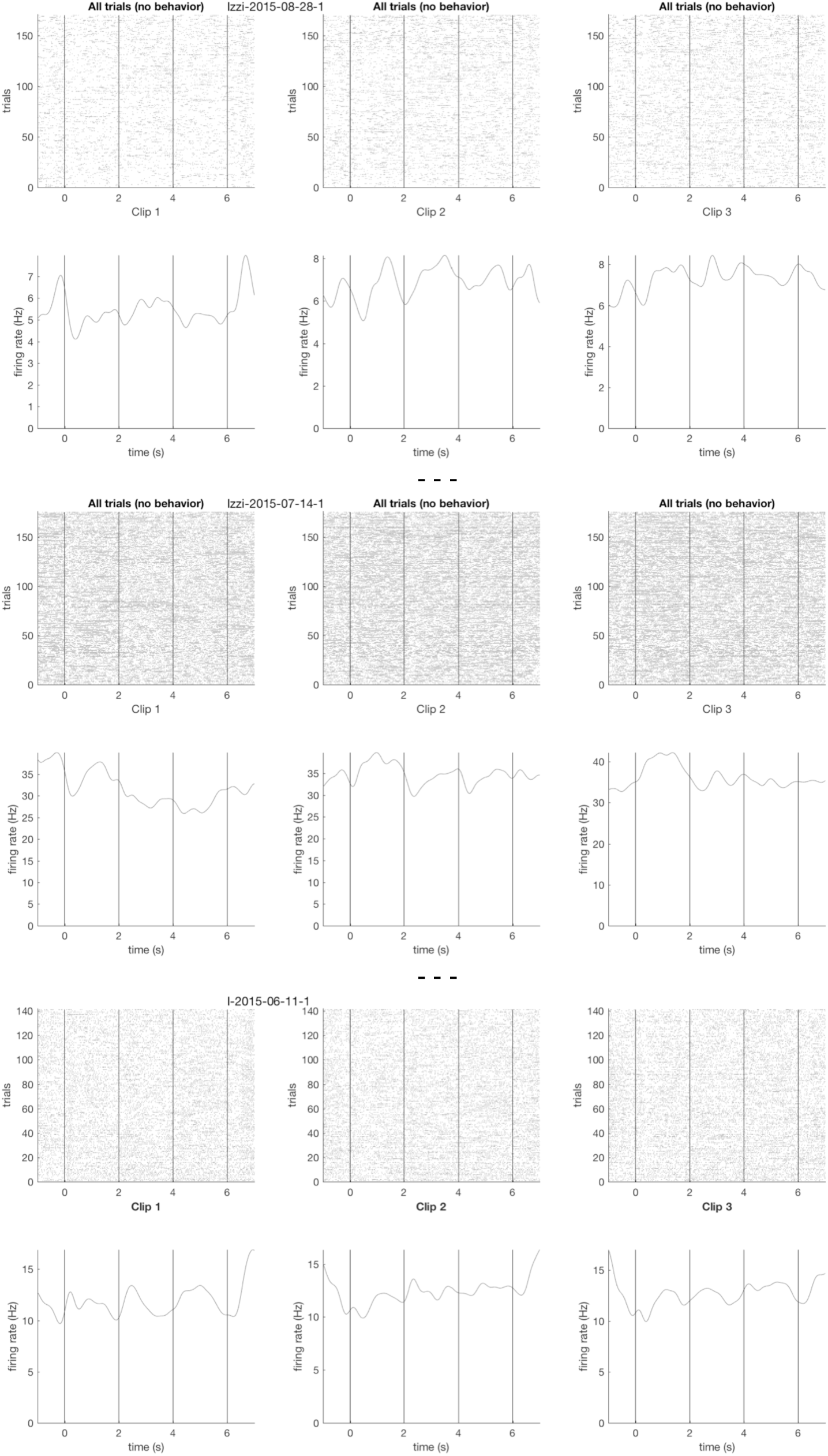
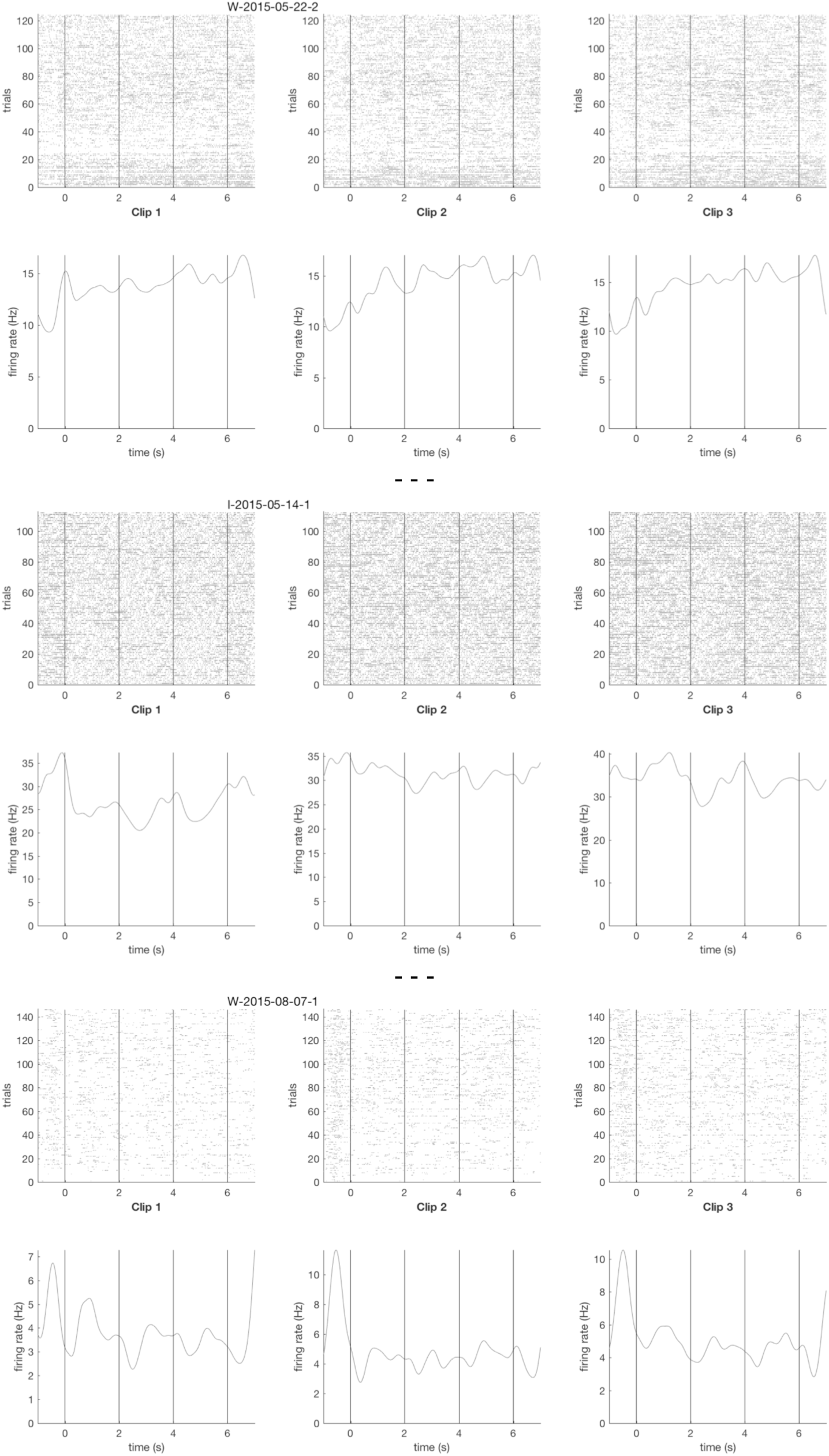
Rasters and PDFs of clips 1-3 for six example neurons in Fig. 4 and Fig. S2.

